# Community-based peptidomic clustering reveals pathogen-specific proteolytic signatures in wounds

**DOI:** 10.1101/2023.12.28.573527

**Authors:** Erik Hartman, Fredrik Forsberg, Sven Kjellström, Jitka Petrlova, Congyu Luo, Aaron Scott, Manoj Puthia, Johan Malmström, Artur Schmidtchen

## Abstract

Recent advances in mass spectrometry-based peptidomics has catalyzed the identification and quantification of thousands endogenous peptides across diverse biological systems. However, the large theoretical peptidomic landscape and high proportion of missing values poses several challenges for downstream analyses and limits the comparability of clinical samples. Here, we present a generalizable computational workflow that clusters peptides with overlapping sequences to reduce the dimensionality of peptidomic data, improve the definition of protease cut-sites, enhance inter-sample comparability, and enable the implementation of reliable and large-scale data analysis methods akin to those employed in other omics fields. We showcase the algorithm by performing large-scale quantitative analysis of wound fluid peptidomes of highly defined porcine wounds and human clinical non-healing wounds. The analysis revealed signature phenotype-specific peptide regions reflecting pathogen-specific proteolytic activity at the earliest stages of colonization, resulting in novel class of potential peptide cluster-based biomarkers.

## Introduction

Peptides are short sequences of amino acids which, like proteins, play diverse and crucial roles in many biological processes. Recent advances in mass spectrometry instrumentation, sample preparation techniques, and data analysis strategies have catalyzed the large-scale studies of peptidomes. While a fraction of endogenous peptides are synthesized *de novo*, a vast peptidomic landscape emerges through the degradation of proteins by endo- and exoproteases (1,2). The interplay between these classes of proteases results in clusters of peptides with overlapping sequences centered around endoprotease cut-sites, introducing variation and redundancy into peptidomic data. The large number of potential peptides combined with the stochastic data collection associated with data-dependent acquisition mass spectrometry makes peptidomes highly dynamic, typically resulting in diverse peptidomes with a low degree of overlap between samples.

Current peptidomic data analysis strategies largely rely on filtering-methods to sift through the large datasets and remove unwanted degradation products to eventually identify some of the relevant bioactive peptides of importance (3–5). This can be accomplished by e.g. predicting the functions and properties of peptides and thereafter selecting the highest-scoring peptides according to a desired metric. While this ‘needle-in-a-haystack’ perspective has proven successful in identifying endogenous bioactive peptides and peptide biomarkers across diverse contexts such as diabetes, organ failure, inflammation, infection, cancer, neurodegenerative disease, and as neuronal peptide hormones (3,4,6–11), it entails removing a substantial fraction of the peptidome, potentially discarding important information about the biological system in question.

Wound infections constitute a substantial societal burden due to their impact on public health and healthcare resources, the challenges involved in their diagnosis, and the pathogens’ resilience to treatments. Several bacterial species, such as *Pseudomonas aeruginosa*, *Escherichia coli*, *Acinetobacter baumanii*, *Corynebacterium*, *Enterococcus faecalis*, and *Staphylococcus aureus* are particularly common in wound infections. Of these, *P. aeruginosa* and *S. aureus* are the most common culprits in burn wounds and surgical site wounds (12) and are on the Global Priority List released by the World Health Organization due to their threat towards human health and their increasing resistance towards antibiotics (13–15).

Bacteria release proteases and other factors modulate the hosts’ proteolytic activity to shape the peptidome landscape, facilitate immune response subversion, and promote bacterial invasion, making wound infections highly proteolytic environments (2,16–20). Concurrently, the host has developed elaborate peptide-based defense systems which aid in combating the pathogen as a part of the innate defense (21,22). The peptidome lies at the interface of this interaction as it reflects the combination of substrate availability, protease landscape, protease activity, post-translational modifications, and conformally exposed cut-sites, which may vary depending on the microenvironment and type of pathogen. This intricate interplay underscores the need to investigate protein degradation and the resulting peptidome for mechanistic insights and as a source for both biomarkers and therapeutic targets for wound infections (2). Large-scale peptidomic studies could potentially provide a more comprehensive perspective on these mechanisms, however, there remains an unmet need for computational methods that capture the entirety of the peptidome.

We present a computational workflow leveraging the inherent clustering of peptides to simplify peptidomic data analysis. By employing a community-based algorithm, we capture the natural clustering of peptides into entities denoted as peptide clusters. The clustering of peptides into functional entities enable the implementation of reliable and large-scale analytical methods akin to those employed in other omics fields. This reduction in dataset dimensionality enhances inter-sample comparability by alleviating the problem of missing data points. Further, it allows for the detection of phenotype-specific peptide regions, shedding light on the impact of proteases, modulating factors, and peptides in the battle between pathogen and host. We apply this method to deconvolute and study the peptidomes of wound fluids from porcine wounds fluids infected by *S. aureus* and *P. aeruginosa*, individually as well as in superinfections, to uncover the infected wound fluid peptidome and find pathogen-specific biomarkers during the earliest stages of bacterial colonization. We then demonstrate that the method generalizes to complex clinical non-healing wounds in human patients. The methodology underlying the creation and analysis of peptide clusters has been provided in a Python package available under an MIT license: https://github.com/ErikHartman/pepnets

## Results

In clinically infected wounds, the bacterial composition depends on the patients’ microbiome as well as other factors, such as pre-existing conditions and patient genotype (23). The time of initial infection is often unknown, and the composition of bacterial species vary, complicating investigations of bacterial infection dynamics on the peptidome. To mitigate these challenges, we used wound fluids from porcine wounds infected with *S. aureus* (N=21) and *P. aeruginosa* (N=17) on day 0. Four of the *S. aureus*-infected wounds were infected with *P. aeruginosa* on day 1, resulting in a double infection. 13 control wounds were not infected. The wounds were covered with a dressing that absorbed the wound fluid, which was changed every 24 hours and analyzed over a time-course of 2-3 days from infection (Fig. 1a) (24). Proteins were separated from the peptidome using molecular weight cut-off filters of 30 kDa, whereby the wound-derived peptides were identified and quantified by tandem mass spectrometry (MS/MS) (25). In total, 15268 peptides were identified across all samples, of which 4557 are unique to a single sample, demonstrating the relatively low overlap between samples. On average, each sample contained 89.5 ± 7% missing values. Most peptides were identified in *P. aeruginosa*-infected wounds and the fewest were in the control wounds (Supplementary Fig. 1). A characterization of the wound fluid peptidomes can be found in Supplementary Notes S1.

**Fig. 1.**
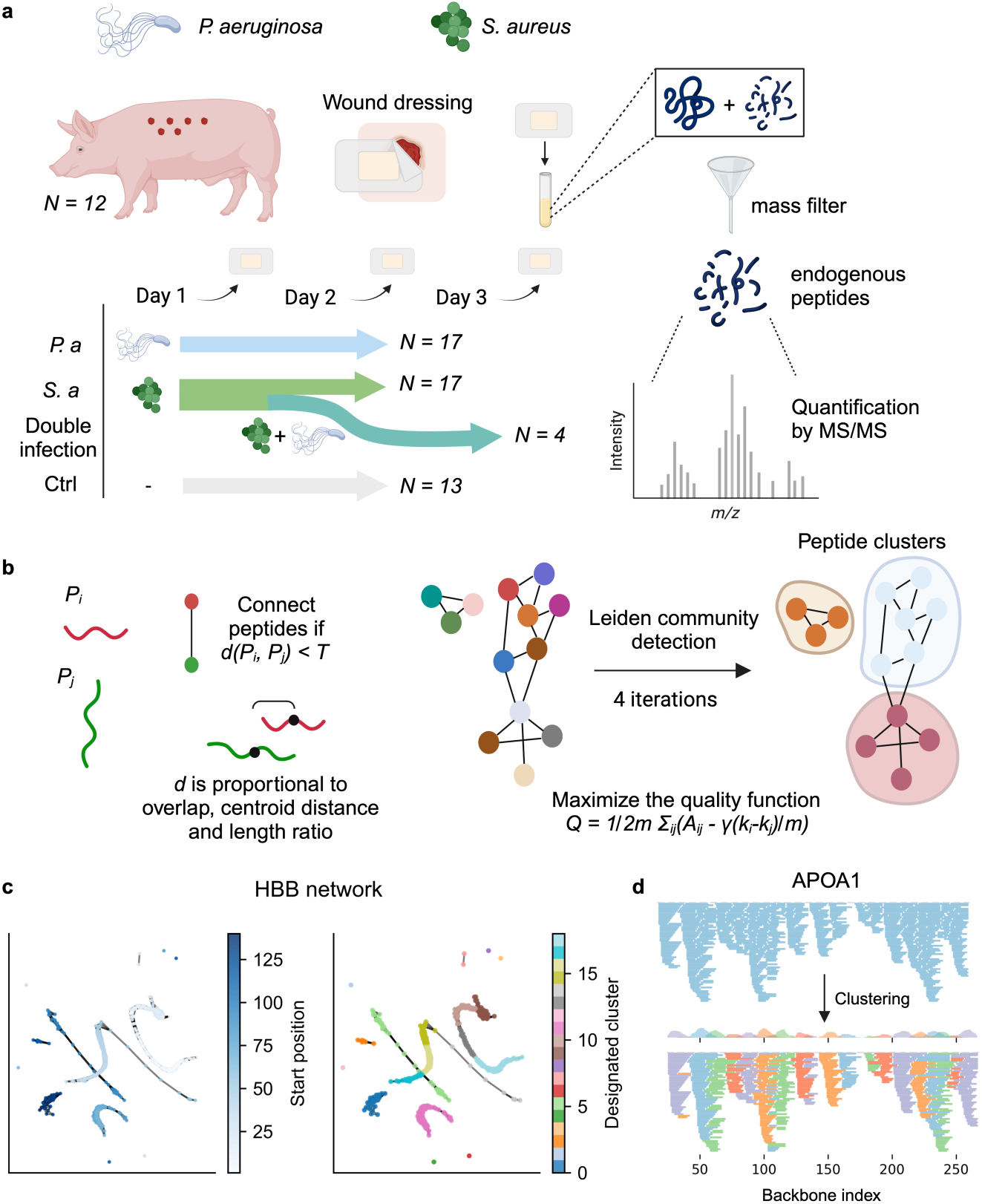
Generating wound fluid peptidomic clusters. **a** Wounds were created on pigs and overlaid with a dressing. The wound drainage, containing proteins and peptides, is absorbed into the dressing. The dressings were changed and sampled every day over a 2-3-day period. The fluid was extracted from the dressings, whereafter proteins were filtered out by applying a mass filter of 30 kDa. The endogenous peptides were subsequently identified and quantified by MS/MS. In total, 21 wounds were infected with *S. aureus*, and 17 with *P. aeruginosa* on day 0. Additionally, 4 of the *S. aureus*-infected wounds were infected with *P. aeruginosa* on day 1, resulting in double infected samples, which were sampled over day 2 and 3. 13 samples were not infected and were used as a control. **b** The initial step in generating peptide clusters entails generating peptide networks (undirected graphs). Peptides are connected if they pass the threshold *d*(*P*_*i*_, *P*_*j*_) < *T* where *d* is a function of peptide overlap, length ratios, and centroid distance, and *T* is a chosen threshold. Here, an optimal threshold of 4 was identified empirically and applied to generate the networks. The resulting network is further partitioned by applying the Leiden community detection algorithm (29) which seeks to maximize the modularity of the network, *Q*, to finally create peptide clusters. **c** Example of the peptide network for HBB where each node represents a peptide. Nodes are colored by starting position (left) and designated cluster after applying the Leiden algorithm (right). **d** Visualization of the clustering algorithm when applied to APOA1. The upper panel shows all the peptides without clustering projected onto the protein backbone. The lower panel shows the results after clustering, where peptides from each cluster are colored differently from its neighbors and depending on what cluster they belong to.

Closer inspection of the peptidome revealed peptide clusters comprised of peptides with partially overlapping sequences, differing by single terminal amino acids, compatible with the influence of exopeptidases. These variants introduce redundancy in the dataset and complicates inter-sample comparisons. Although methodologies aiming to cluster linear peptide sequences have previously been developed to reduce the complexity of epitope-related data (26–28), they do not take peptide length and proximity in the protein backbone into account, which is crucial to the clustering of the peptidome. Therefore, we developed an algorithm that clusters similar peptide sequences by their similarity and proximity in the protein backbone. Therefore, we developed an algorithm that clusters similar peptide sequences with respect to their proximity on the protein backbone.

To initialize clustering, protein centered peptide networks are generated for each protein separately by connecting peptides with overlapping amino acid sequences, similar lengths, and a centroid distance below a certain threshold (Fig. 1b, equations 1-5 in Methods). The topology of the resulting networks depends on the peptide content and may result in small islands of distinct and highly separated clusters, or large connected components as a result of continuous overlap between peptides derived from different cut-sites. To separate the connected components, the resulting networks were further partitioned by applying the Leiden community detection algorithm (29), seeking to maximize the modularity of the network (Fig. 1b). This partitions the large connected components into highly connected subcomponents, which effectively separates between the unique clusters. An example of the network for hemoglobin subunit beta (HBB) and the resulting designated clusters can be seen in Fig. 1c. Another visualization of the clustering when projecting the peptides in APOA1 to the protein backbone can be seen in Fig. 1d. Peptides within a cluster have consistent biophysical properties, motivating the use of peptide clusters as a functional entity (Supplementary Notes S2).

### Characterization of wound fluid peptidomes

To characterize the wound fluid peptidomes, we apply our community-based clustering strategy to the complete dataset of 15268 unique peptide sequences, resulting in 761 clusters, thereby reducing dimensionality by 95%. The average number of missing values was reduced by 70.5 ± 13.3%, representing an increase in present values by 320 ± 87%. The clusters were quantified by taking the sum of the three most abundant peptides (topN-method). Grouping the peptide clusters using hierarchical clustering shows two distinct groups. One group predominately contains shared peptide clusters found in most of the samples whereas the other group contains mostly peptide clusters unique to a smaller set of samples (Fig. 2a). Dimensionality reduction by UMAP showed that the quantified clusters separate the sample types into distinct groups (Fig. 2b). In general, peptide clusters are more abundant and numerous in infected wounds, both in terms of intensity and number of unique clusters (Fig. 2c), indicating higher proteolytic activity during infection. The *P. aeruginosa*-infected wounds contain more abundant and unique clusters compared to those infected by *S. aureus*, while the double infected wounds show similar cluster intensities as wounds infected by *P. aeruginosa*. The clustering enables statistical comparisons between sample types using the DE-score (30). In total, 98 clusters had a *log*_2_(fold change) > 4 and a p-value < 0.01, demonstrating that the peptide clustering strategy supports identification of clusters that are differentially abundant between sample types. In Fig. 2d, the abundances of the 5 top-scoring clusters in *S. aureus* and *P. aeruginosa* respectively, are shown.

**Fig. 2.**
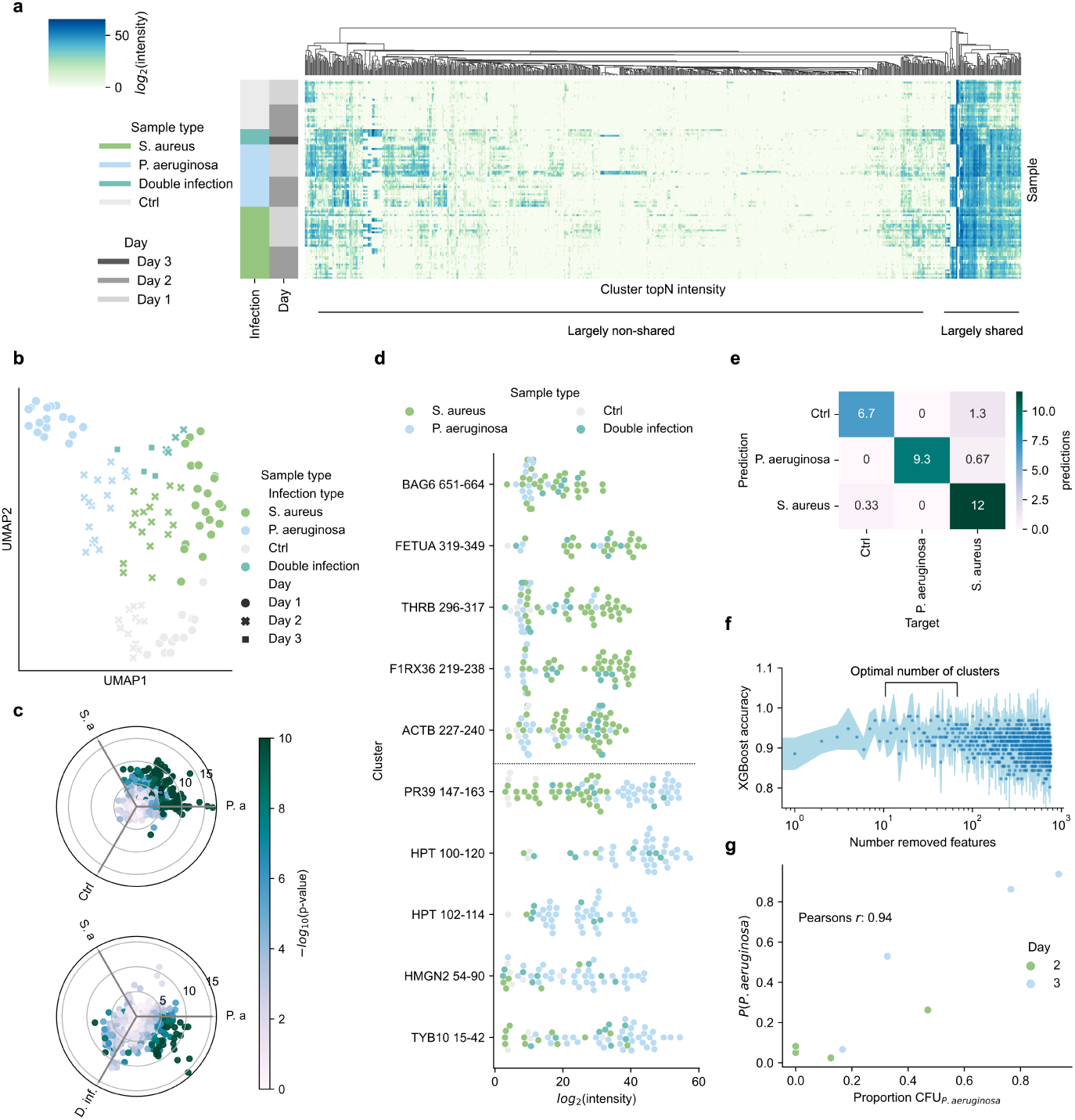
Peptide clusters enable large scale analysis. **a** Clustermap of peptide clusters. The color intensity is proportional to the cluster quantity calculated by the topN method on the *log*_2_-transformed intensity. Hierarchical clustering is performed on the peptide clusters (columns). **b** Uniform manifold projection (UMAP) of the quantified clusters colored by sample type. The shape indicates the day the samples were taken. **c** A polar projection where each scatter represents a cluster. The position of the cluster is computed as ∑ **^v**_*i*_ where *i* are the sample types |**v**_*i*_| the cluster quantity. Each sample type is separated by 120°. The color intensity is proportional to the −*log*_10_(*p*−*value*) as determined by ANOVA, clipped to ≤ 10. **d** Swarmplot of the *log*_2_-intensities of the top 5 clusters by DE-score for *S. aureus* (upper) and *P. aeruginosa* lower. **e** Average confusion matrix from k-fold cross-validation (*k* = 3) of an XGBoost classifier. **f** Recursive feature elimination using SHAP to estimate feature importance was performed to investigate the number of optimal features for an XGBoost classifier. The mean accuracy after removing a given number of features is shown as scatters. Error bands represent ±1 SD. An optimal number of features lie in the range of 10-60 features (out of 781 total). **g** A support vector machine (SVM) with a linear kernel was trained to classify bacterial species of infection based on the quantified peptide clusters. The y-axis shows the prediction probability for *P. aeruginosa* infection when given the double infected samples, and the x-axis shows the proportion of CFU for *P. aeruginosa* compared to *S. aureus* for the samples. The correlation between SVM probability and *P. aeruginosa* CFU-proportion is 0.92 (Pearson’s r).

An important property of the reduced data is that standard omics-esque machine learning-based classification of samples can be performed. Here, we trained an XGBoost (31) classifier to distinguish between the controls, *S. aureus* and *P. aeruginosa*-infected samples, and achieved a classification accuracy of 92 ± 1% (Fig. 2e). To investigate how many clusters are indicative of infection, a modified version of recursive feature elimination using SHAP (32) was applied to iteratively remove features deemed less important to the classifier. Optimal performance was achieved when utilizing 10-60 clusters (Fig. 2f). The top 50 clusters identified as the most important ones are listed in Supplementary Table 1.

Infections are not binary but exist on a continuum, as some wounds are more heavily colonized than others. We hypothesized that the amount of colonized bacteria was reflected in the peptidome, and that the output probability of a classifier trained on the peptidomic content therefore would be indicative of the level of colonization. Similarly, the classifier probability in a double infection would reflect the fraction of colonization by the respective species. To test if the classifier can predict the CFU-composition in the double infected wounds, we trained an SVM to distinguish between the single-infected samples, and then determined the probability of *P. aeruginosa* infection in the double infected samples. The correlation between the fraction of CFU_*P.aeruginosa*_ and output probability was 0.94 (Pearson’s r) (Fig. 2g). This demonstrates that the level of bacterial colonization is reflected in the peptidome, indicating that the effects of either specific bacterial proteases or modulating factors are captured in peptide clusters.

### Clusters can be utilized to identify bacterial protease activity

To identify the protease activity which has given rise to the different peptidomes, we first analyzed the wound fluids using zymograms. Here, three distinct patterns were seen in control and single infections, where *P. aeruginosa*-infected wounds showed enzymatic activity with degradation in the 75-50 kDa and 100 kDa-regions (Fig. 3a). The *P. aeruginosa*-related enzymatic activity was not observed until day 3 in the superinfected samples, suggesting that *P. aeruginosa*-colonization is delayed when there is a pre-existing *S. aureus*-colonization (Fig. 3b). The protein content was investigated with SDS-PAGE, showing little to no difference between sample types (Supplementary Fig. 3).

**Fig. 3.**
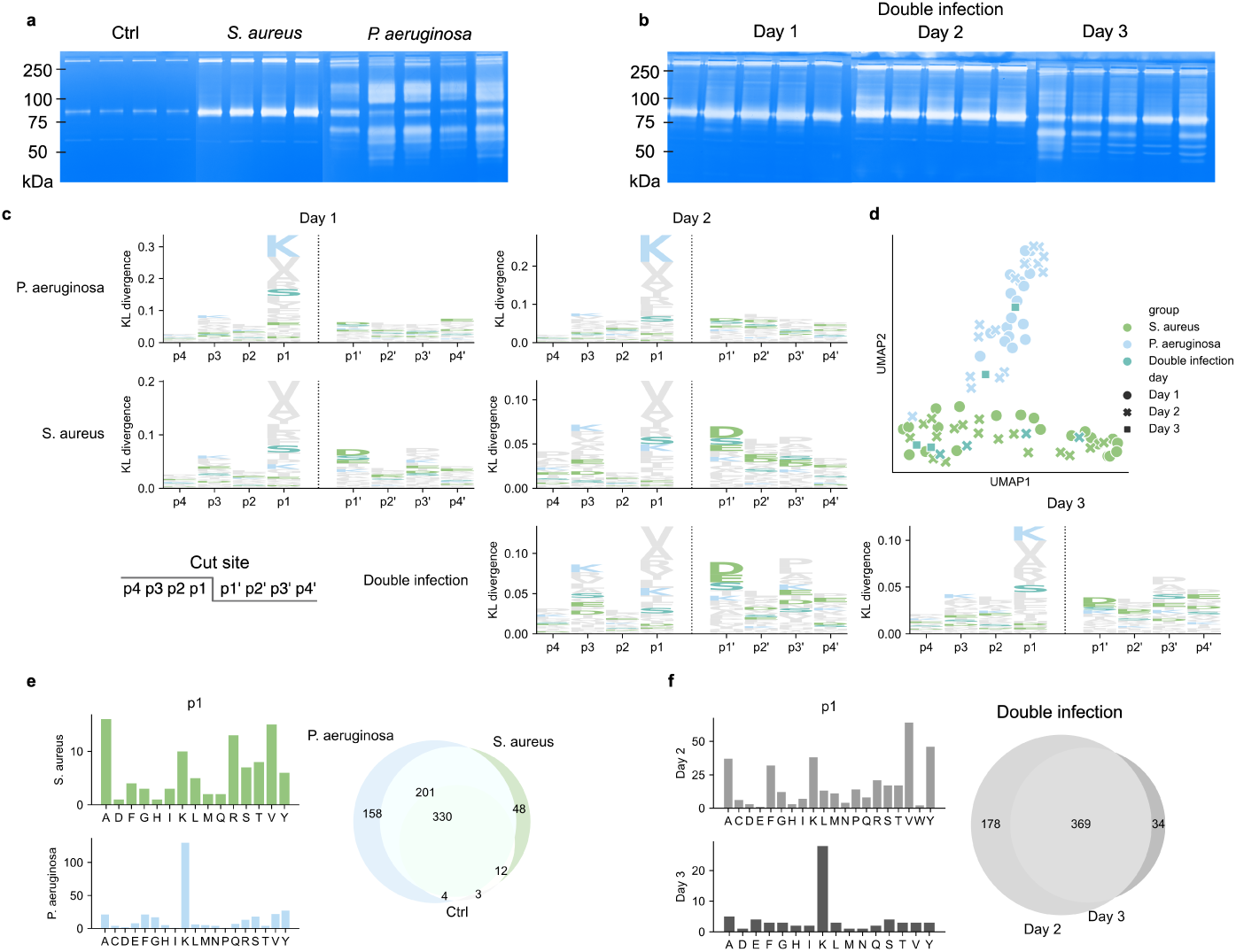
Identification of proteases utilizing peptide clusters. **a,b** Zymograms for the single infections and control during day 1 (**a**) and for double infected samples during the timespan (**b**). Double infection on day 1 is a single *S. aureus* infection. An approximate kDa ladder is shown to the left of the gels. **c** Logoplots of cut-site specificity (p4-p4’). The height of the column is calculated by the Kullback-Leibler divergence between the weighted amino acid distributions for the sample types against control. The amino acid distributions are generated by identifying the amino acid at the most commonly occurring terminals for each peptide cluster, as these are the most likely endoprotease cut-sites. **d** Projection of the KL-weighted p4-p4’ amino acid frequency to two dimensions using UMAP. **e** The left panels show the p1 amino acid distributions of the cut-sites for the unique clusters in *S. aureus*-infected wounds and *P. aeruginosa*-infected wounds. The right panel shows a venn-diagram of the peptide cluster overlap. **f** Similarly to in **e**, but for the double infected samples.

Investigating the p4-p4’ regions surrounding peptide terminals can reveal specific proteolytic activity in wound infections, as proteases typically exhibit specificity in this residue window. Each cluster is associated with two cut-sites (for non-terminal peptides) defined by the most common terminal position in each cluster and sample. The amino acid distributions in the p4-p4’ windows were weighted by the topN-intensity, generating a weighted amino acid distribution for each sample type at each position surrounding the cut-site. The distributions of infected sample types were then compared to the control-distribution using the Kullback-Leibler (KL) divergence. Across all infected sample types, the largest divergence was observed in the p1-position, indicating a different protease-profile compared to the control. The *P. aeruginosa* infected samples have a cut-site specificity towards lysine at the p1-position, whereas *S. aureus* infected wounds have a specificity towards valine and arginine. In *S. aureus*- infections, there is also a specificty at the p3, p1’ and p3’ positions, although not as distinct as the one at p1. The superinfected wounds exhibited an *S. aureus*-like profile on day 2, whereafter the typical *P. aeruginosa* lysine-specificity emerged on day 3 (Fig. 3c) which correlates with the patterns in the zymograms. This reiterates that *P. aeruginosa*-colonization is delayed when there is a pre-existing *S. aureus*-colonization (Fig. 2b) and that the bacteria-specific proteolytic activity can be discerned in the cut-sites. Projecting the KL-weighted cut-site amino acid frequencies to two dimensions using UMAP, separates the samples cluster based on infection, but not on day, in contrary to what was observed when projecting the quantified clusters (Fig. 3d). In total, 158 peptide clusters are uniquely identified in the *P. aeruginosa*-infected wounds over the complete time span. These clusters are almost entirely enclosed by cut-sites with a lysine at the p1-position (Fig. 3e). A similar profile emerges on day 3 in superinfected samples (Fig. 3f). These results show that the type of clusters, cluster sizes, and intensity changes in a time-dependent fashion, whereas the cleavage patterns are largely similar over time and dependent on the type of bacterial pathogen.

### Identifying pathogen-specific peptidomic clusters in HMGB1, HPT and PR-39

The previous analyses have demonstrated that the algorithm enabled identification of differentially abundant clusters and evidence of bacteria-specific proteolytic activity in the peptidomic content of infected wound fluids. In Fig. 4 we showcase 3 proteins that exemplify different degradation patterns dependent on infection type and which potentially could be used as pathogen-specific biomarkers.

**Fig. 4.**
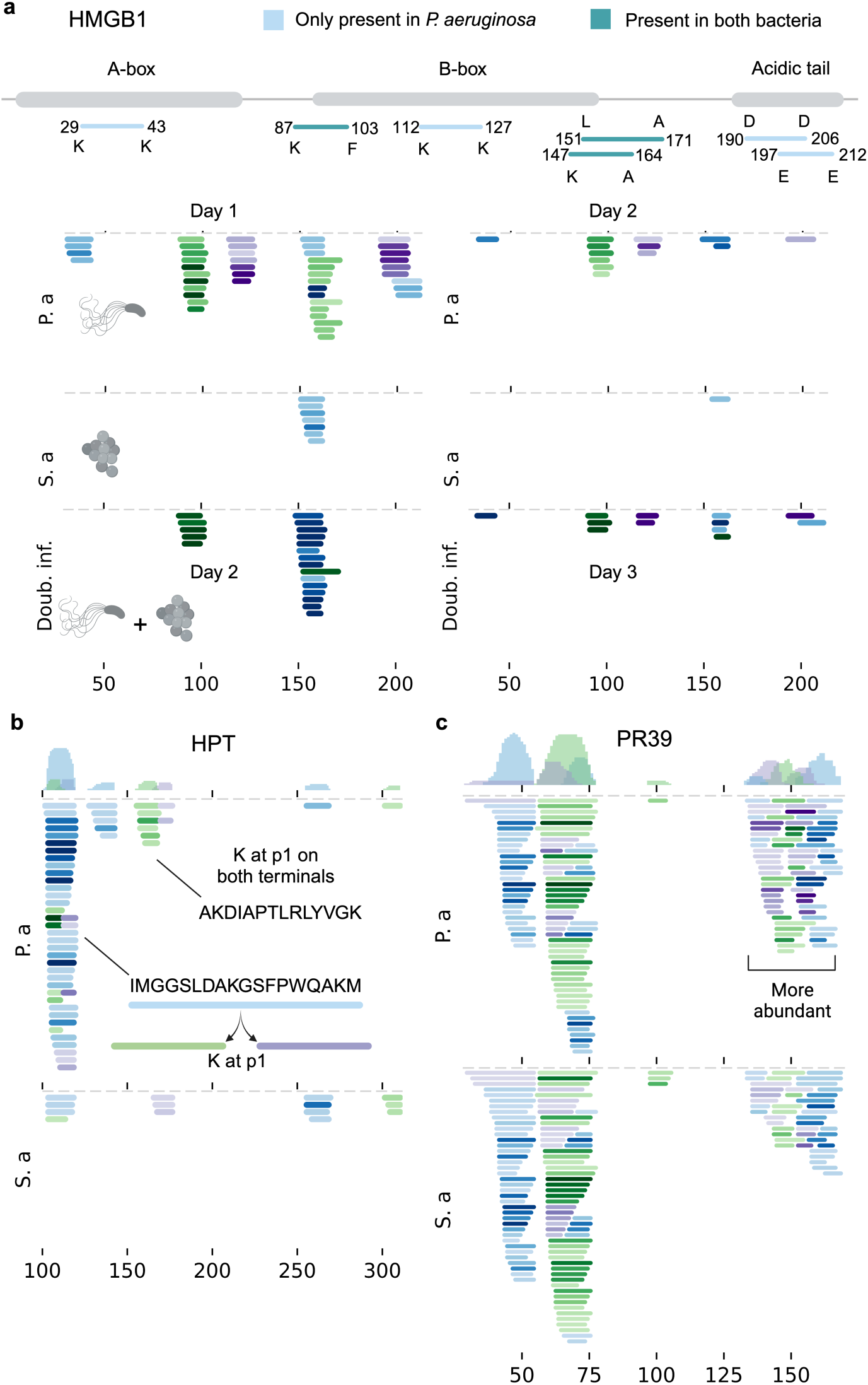
Identification of potential pathogen-specific biomarker clusters. Visualization of the peptidome projected onto proteins. In all figures, the top panel contains a histogram showcasing the number of identified residues at a given position, separated on peptide clusters. The clusters are colored repeatedly in green, blue and purple. The color intensity is proportional to the mean peptide intensity. **a** The cleavage pattern of High-mobility group box 1 (HMGB1) for the single and superinfected samples. In **b, c** the samples for single infection over day 1 & 2 are pooled. **b** Haptoglobin (HPT) contains differentially abundant and unique peptide clusters in *P. aeruginosa*-infected wounds. A major cluster is cleaved into two subpeptides only present in *P. aeruginosa*-infected wounds. **c** Peptides in the antimicrobial part of PR-39 (131–169) are more abundant in *P. aeruginosa*-infected wounds.

High mobility group-box 1 (HMGB1) exhibits different peptide profiles in *P. aeruginosa*-infected wounds and *S. aureus*-infected wounds. Cleavage of HMGB1 in wounds infected by *P. aeruginosa* produced peptides which were not identified in *S. aureus* infections or control. These clusters appeared in double infected wounds on day 3, when the bacterial load of *P. aeruginosa* had increased (Fig. 4a). Haptoglobin (HPT) also exhibits a different peptidomic profile in *P. aeruginosa* infections. Peptides from the region 100-120 are cleaved into two sub-peptides which are identified in *P. aeruginosa*-infected wounds (Fig. 4b).

In certain proteins, the degradation patterns are highly similar, but certain clusters are differentially abundant. Amongst these proteins is the antimicrobial protein PR-39. Several clusters from the antibacterial region of PR-39 (131–169) are identified in both *P. aeruginosa* and *S. aureus*-infected wounds (Fig. 4d). All clusters of PR-39 exhibit similar abundance, except for clusters in this region, which are more abundant in *P. aeruginosa*-infected wounds (Supplementary Fig. 5 & 6). Further discussion of the biological significance of the exhibited protein degradation can be found in Supplementary Notes S3.

### Characterizing contaminated wounds with peptide clusters

During bacterial analysis of the wounds, it was noticed that 4 of the wounds which were infected with *S. aureus* on day 0 were contaminated prior to inoculation (24). The enzymatic activity was therefore analyzed with zymograms, showing similarities to patterns typical for *P. aeruginosa* on day 1, and *S. aureus* on day 2 (Fig. 5a). The peptidome of these wounds were analyzed in means of finding out the peptidomic nature of the double infection. Using the same machine learning classifier which was used to estimate the CFU-composition previously, the probability of *P. aeruginosa* infection was determined in the unintentionally infected samples. The samples were given a high probability of being *P. aeruginosa*-infected on day 1, which then shifted to *S. aureus* on day 2 (Fig. 5b). In a UMAP-projection, the samples on day 1 cluster with *P. aeruginosa* samples, whereafter they shift to *S. aureus* (Fig. 5d).

**Fig. 5.**
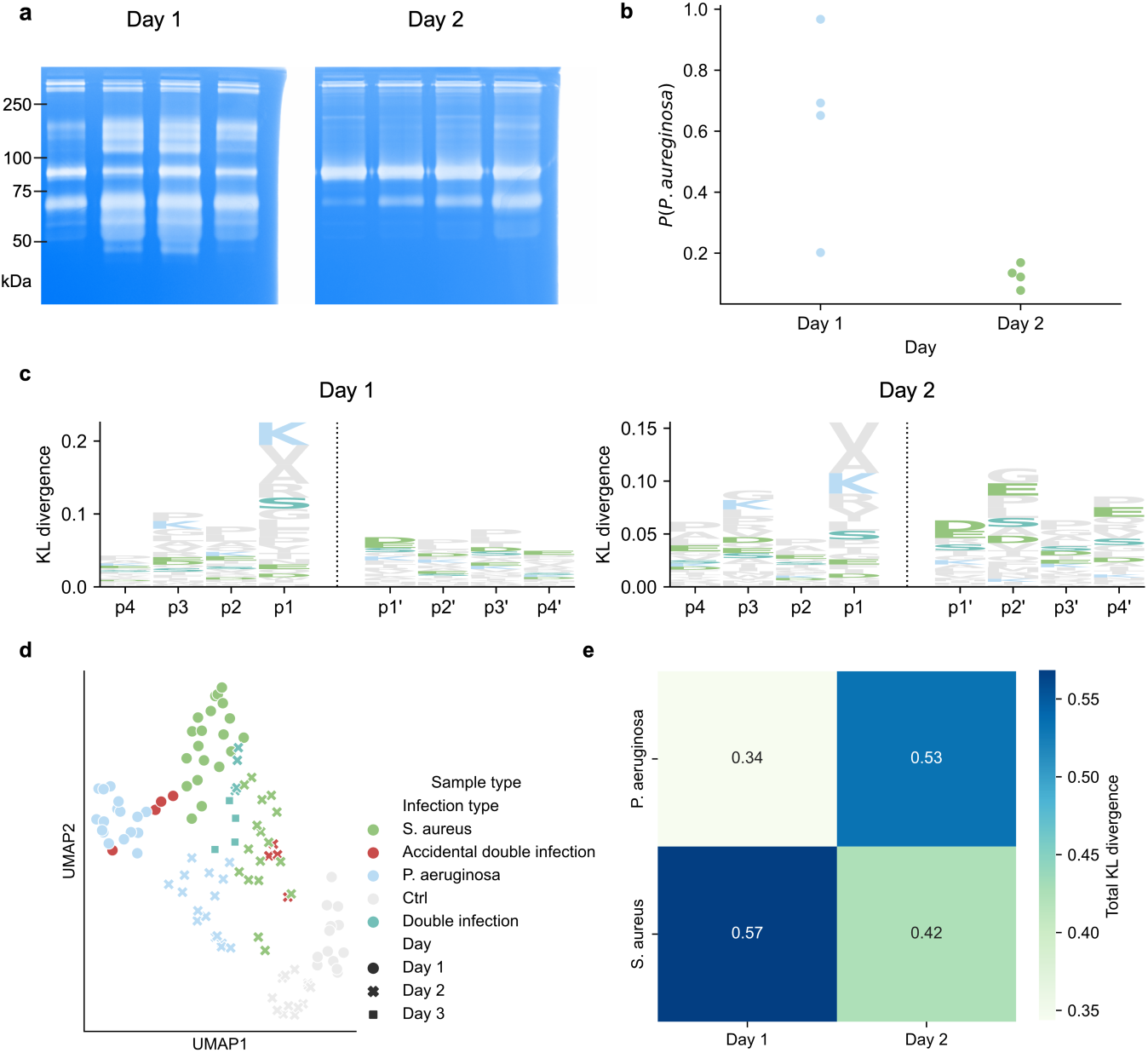
Characterisation of contaminated wounds. **a** Zymograms for the accidentally double infected wounds. An approximate kDa ladder is shown to the left of the gels. **b** The classification probability given by an SVM of the sample being *P. aeruginosa* for the accidentally double infected samples. **c** Logoplots of cut-site specificity (p4-p4’) for the accidental double infections. The height of the column is calculated by the Kullback-Leibler divergence between the weighted amino acid distributions for the sample types against control. **d** Uniform manifold projection (UMAP) of the peptide intensities colored by sample type and shaped determined by the day the sample was taken. **e** Heatmap of the total KL divergence between the amino acid distribution of accidentally double infected wounds (with days as columns) and the single infections (pooled over day 1 and 2). The total KL divergence against *P. aeruginosa* increases between day 1 and 2, meaning that the cut-site specificities get increasingly dissimlar.

On day 1, the cut-site specificity resembles that of *P. aeruginosa* or the double infection on day 3, as lysine is the most common amino acid at p1. This changes on day 2, where the cut-sites resembles *S. aureus* or double infection(Fig. 5c). The cut-site specificity was quantified by taking the sum of the KL-divergence in the p4-p4’ window, showing that the divergence to *S. aureus* decreases from day 1 to day 2, while there is a slight increase against *P. aeruginosa*. Together, this demonstrates the utility of the methodology by identifying subtle differences in proteolytic patterns.

### Characterisation of wound fluid from non-healing wounds

Lastly, we sought to investigate whether the methodology developed here generalizes to human samples by applying it to human wound fluids from patients with non-healing wounds. Common bacteria found in non-healing wounds include *S. aureus*, *P. aeruginosa* and *Enterococcus* species (33). A total of 18 samples were analyzed with the MS/MS workflow described above, resulting in 41741 peptides. Out of these, 10 samples were characterized as primarily being colonized by *P. aeruginosa*, and 8 by being primarily colonized by *S. aureus*. The qualitative bacterial composition of the wounds was determined using MALDI, and the protease activity was analyzed with zymograms (Fig. 6a). The complete list of identified bacterial species and sample specifications can be seen in Supplementary Table 2 and described in Supplementary Notes 4.

**Fig. 6.**
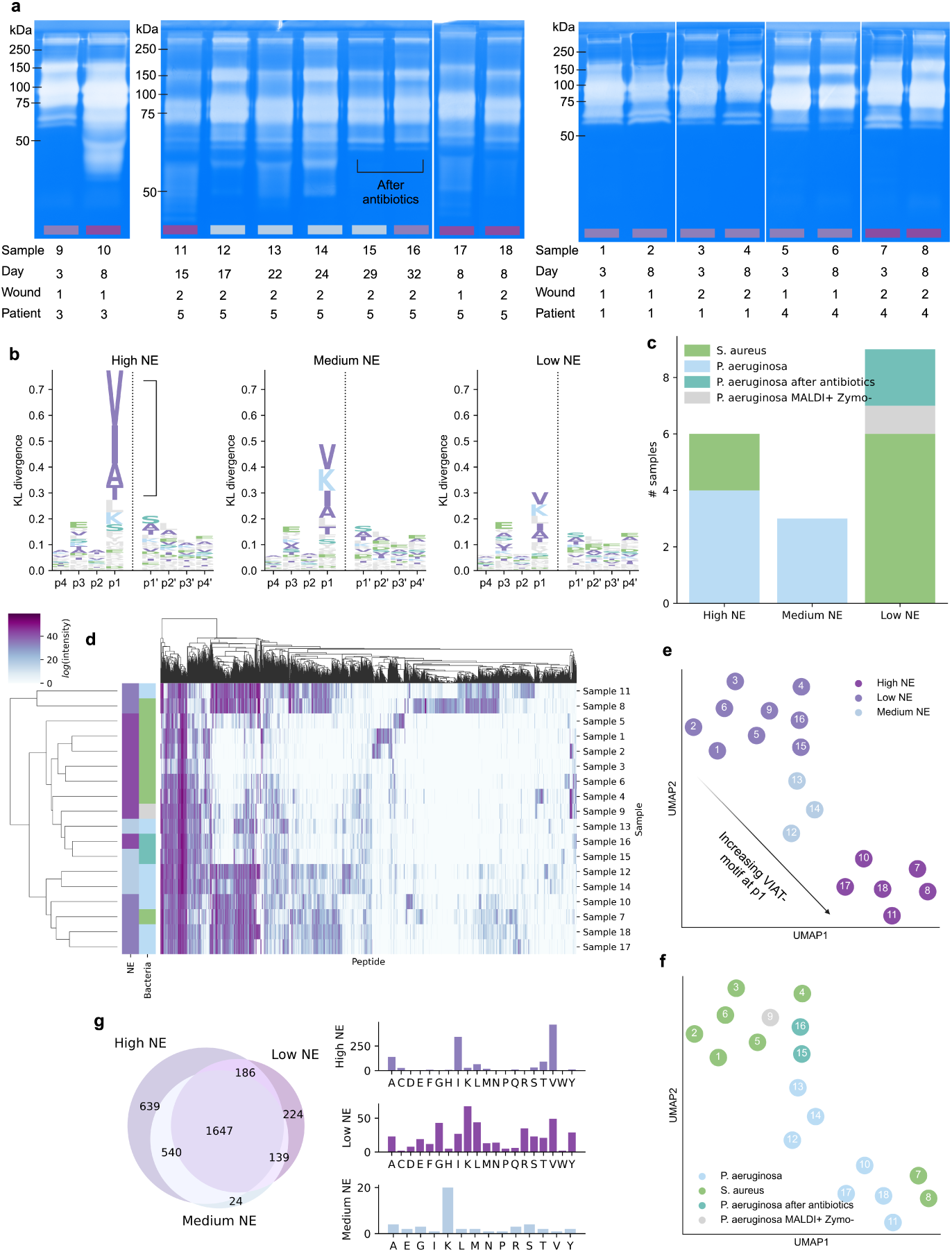
Characterisation of human chronic wound fluid peptidomes. **a** Zymograms of all 18 samples. The bar under the lane is colored based on the level of NE-like cut-site specificity. A kDa ladder is shown to the left of the gels. The gels for sample 9 and 10 were shorter and therefore have their own reference ladder. **b** Logoplots of the mean p4-p4’ amino acid frequency for three groups of samples identified by clustering, which correspond to different levels of NE-like specificty at the p1-position. In addition to previous colorations, valine, isoleucine, alanine and threonine are now colored purple. **c** The number of samples with different bacterial species determined by MALDI and zymograms with regards to the different level of NE-like specificity. **d** Clustermap of log-transformed peptide cluster intensities. The row colors correspond to the identified groups in **b** and on the bacterial species identified in the zymograms and MALDI. **e** UMAP of the p4-p4’ amino acid frequency for all samples, colored by identified clusters which correspond to level of NE-like specificity at the p1-position. **f** UMAP of the p4-p4’ amino acid frequency for all samples, colored by the consensus between the results from MALDI and the zymograms. **g** The amino acids identified at the p1-position for the unique clusters when comparing samples with different levels of NE-like specificity.

Peptide clusters were generated with the same settings as previously applied to the porcine samples, finally resulting in 3254 clusters - reducing the dataset size by 92% and increasing the fraction of present values by 290 ± 30%. Analyzing the cut-sites revealed a highly specific motif at the p1-position. The motif is identical to that of human neutrophil elastase (NE), with a strong specificity for valine, alanine, isoleucine and threonine, in descending order (34,35) (Fig. 6b). All *P. aeruginosa*-infected wounds revealed a moderate or high level of NE, while a majority of *S. aureus*-infected samples had a low level of NE-like specificity (Fig. 6c). Performing hierarchical clustering on the peptide cluster intensities revealed that they largely cluster on NE activity and bacterial species (Fig. 6d, Supplementary Fig. 7a). Dimensionality reduction using UMAP was performed on the KL-weighted p4-p4’ amino acid frequencies, revealing three clusters which can be recognized as having different degrees of NE-like p1 specificity (Fig. 6e). These also show a correspondence with bacterial colonizer and the zymogram appearance, as e.g., sample 9, 15 and 16 cluster with *S. aureus*.

The clusters exclusively identified in the high NE-samples were enclosed by cutsites with valine, alanine and isoleucine at the p1-position. Similarly, the clusters exclusively present in the moderate NE-samples which are mainly colonized by *P. aeruginosa*, showed a specificity against lysine in the p1-position (Fig. 6g) which is not identifiable if stratifying on primary bacterial colonizer (Supplementary Fig. 7b). The cut-sites were analyzed after removing all clusters enclosed by valine, alanine, isoleucine or threonine. Performing UMAP on the resulting KL-weighted p4-p4’ amino acid frequencies reduced the influence of NE and separated the samples based on bacterial colonizer (Supplementary Fig. 8a,b). In conclusion, these results show that the peptide cluster strategy proposed here can enable the identification of the bacterial species of the primary colonizer in complex non-healing human wounds based on the peptidome alone.

## Discussion

Understanding the impact and the underlying processes behind protein degradation has enhanced our understanding of biological systems and represents an unexplored resource for the identification of new biomarkers and therapeutic targets. However, large-scale peptidomics analyses present challenges rooted in the inherent diversity and scale of the peptidomic landscape. To address these challenges, we present a method that deconvolves the peptidome by using networks and community detection algorithms to approximate optimal partitioning. The algorithm results in a data-reduction of 93-95% in our datasets, combating the curse of dimensionality and opening avenues for analytical strategies similar to those successfully employed in other omics fields. The method also enables a new definition of an endoprotease cut-site as the most common terminals to a peptide cluster, leading to improved cut-site analyses. Importantly, we demonstrate that classification models can utilize peptide clusters to distinguish between samples, identify important clusters, and estimate the type and level of pathogen-specific colonization and activity. This method not only advances our understanding of the peptidome but also enhances the precision and interpretability of large-scale peptidomics analyses in complex biological contexts.

The data and algorithm presented here has several shortcomings that warrant consideration. Firstly, the algorithm is dependent on the manual selection of parameters, such as the threshold selected when creating networks and the resolution parameter during partitioning. The values for these parameters were identified empirically by visualizing the data and seeing what made intuitive sense. No data-driven parameter selection algorithm was found to generate optimal clustering, mainly because the optimization goal was difficult to define. Secondly, the datasets presented here are limited, with 111 porcine samples and 18 human samples. Given the complexity and diversity of wounds and the peptidome, there is a need for studies to validate our findings and investigate the wound fluid peptidome further.

Despite these limitations, we identify unique cut-sites dependent on the species of the primary wound colonizer. Generally, cut-sites vary mostly in the p1-position. Wounds infected by *P. aeruginosa* exhibit a large specificity towards lysine residues at p1 which is compatible with the reported specificity of its secreted virulence factor protease IV (36). In human non-healing wounds, we see a large influence of neutrophil elastase-like proteases which is in agreement with previous studies demonstrating elevated elastase levels (37). Although we do not identify any known proteases in the cut-site specificity of *S. aureus*- infected wounds, we identify differentially abundant and unique clusters in both cases. These generally correspond to clusters derived from known inflammatory proteins, such as HMGB1, PR39 and protegrins when infected by *P. aeurginosa*, and cytoskeletal proteins when infected by *S. aureus*.

The methodology presented here has the potential to instantiate a new branch of peptidomics research, in which large-scale data-driven analyses are used to identify peptidomic differences in different biological contexts. Here, we have demonstrated its utility in wound infections, however, we envision similar methodologies to be applied to a plethora of contexts, ranging from infectious diseases to cancer and metabolic disorders.

## Methods

The goal of generating peptide clusters is to group highly similar and proximate peptides into a single entity. This coincides with capturing the endopeptidase activity, while filtering out the effect of exoproteases which generate clusters of highly similar peptides. If all possible variations of peptides were present in the data, this would be an easy task, since all peptides could be merged onto the longest variant. However, due to random processes and sampling stochasticity, the data becomes noisy. Additionally, sometimes a peptide is cleaved into two subpeptides, in which case information is lost when merging onto the parent peptide. These issues call for a fuzzy algorithm to yield the most optimal grouping of peptides. Here, we devise an approach which utilizes networks and community detection to approximate the optimal grouping of peptide clusters.

### Generating peptide clusters

The first step in generating peptide clusters is to generate weighted peptide networks, where peptides we consider being part of the same cluster have a low distance between them. The distance between each pair of peptides is calculated using a distance function, *d*. The distance function devised and applied here takes peptide overlap, centroid distance, and peptide length ratios into account. The rationale behind the choice of these variables are the following: peptides with a high degree of overlap are likely to belong to the same cluster. Long peptides with high overlap, but a large distance between their centroids are unlikely to be from the same cluster. Peptides with different lengths could belong to the same cluster, but should not be connected directly. If intermediate products are present, they will have an indirect connection, however, if not, they should not be connected, as they are most likely from different clusters. The different parts of the distance function for peptides *p*_*i*_ and *p*_*j*_ are:

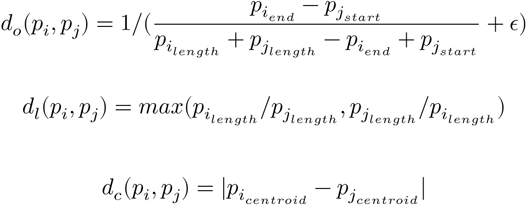

Where *ϵ*is a noise variable of 10^−8^, *p_i_length__* the length of peptide *i* and *p_i_centroid__* the centroid position of peptide *i*. The complete distance is the sum of the factors:

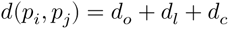

And the peptides are connected if:

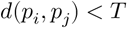

where *T* is an arbitrary threshold which is purpose specific. In our data, an optimal threshold of 4 was identified empirically. A network is created for each protein separately. The created networks are partitioned into isolated components, with weaker connections between clusters with lowly overlapping components. To further partition the network, the Leiden community detection algorithm is applied.

The Leiden community detection algorithm is highly similar to the more common Louvain detection algorithm, but improves on it by guaranteeing well-connected communities (29). These algorithms seek to maximize the modularity of the network, which is calculated as:

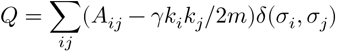

where *A* is the adjacency matrix, *k*_*i*_ is the weighted degree of node *i*, m is the total sum of edge weights and *δ*(*σ*_*i*_, *σ*_*j*_) = 1 if *i* and *j* are in the same community. *Γ* is the resolution parameter which defines the expected number of communities. In our dataset, the optimal resolution factor was investigated empirically and given a value of 0.8.

It was noted that if a protein contained regions of highly varying peptide density, the algorithm tended to split clusters in high density regions even though the peptides overlapped to a great extent. To correct for this, a manual step of merging clusters was applied, where clusters were merged if the cumulative distance of each termini was below given threshold. Here, a threshold of 2 was used. Lastly, clusters with fewer than 3 peptides were removed from the dataset since these could not be quantified with the topN-method when N=3.

### Analyzing peptide clusters

Cluster quantification can be conducted similarly to protein quantification in proteomics research. Here, clusters were quantified by taking the total value of the top 3 most intense peptides as is common in label-free proteomics quantification (38). To investigate what clusters differed the most between the sample types, differential abundance analysis and machine learning based feature extraction was performed. For pair-wise differential abundance analysis, p-values were calculated using students t-test. The most differentially abundant peptide clusters were selected using the DE-score (30). The DE-score is the pythagorean sum of the normalized fold change and p-value, and was modified so that the logarithmized fold change was sign-dependent.

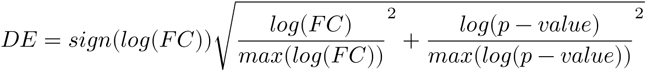

Thereby, the top peptide clusters for both groups included in the analysis can be extracted separately. When comparing more than two groups, p-values were calculated using ANOVA.

To investigate if classification-models could be used to distinguish between sample types, classificaton was performed with an XGBoost classifier. The classifier takes the scaled cluster intensities as input. To investigate how many peptide clusters and which peptide clusters were important for classification, a modified recursive feature extraction scheme was implemented. Here, features are iteratively removed based on their SHAP value, as features with low SHAP are considered unimportant for the classifier.

It was hypothesized that a classification model could be used to estimate the fraction of pathogen colonization in superinfected wounds. An SVM with a linear kernel was trained to classify the single infections, whereafter it was used to classify the superinfected wounds. An SVM with a linear kernel was chosen so that the decision boundrary and thereby the output probability would reflect the prediction certainty in a linear fashion, so that it could be compared with the CFU-fractions for the different bacteria. The output probabilities were correlated with the CFU-fractions using Pearson’s r.

The most probable cut-site giving rise to each cluster is the most common terminal, i.e., the mode of the terminal positions for each cluster and sample. To identify cut-site specificity, windows spanning 8 amino acids surrounding these sites were considered (p4-p4’). The amino acid distribution for each sample type and position was weighted with the mean peptide cluster intensity. The influence of the pathogens on the cut-sites was compared against control. To quantify the difference between these distributions, the Kullenback-Leibler divergence was computed as follows:

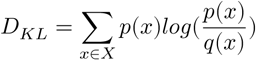

Where *p*(*x*) is the amino acid distribution for the infected sample type and *q*(*x*) for the control. The divergence was used to weigh the amino acid frequency at each position, before displaying them in a logo-plot.

The cut-site amino acid frequency distribution is a matrix of dimensions 23 × 8. Stacking this matrix into a vector of length 23 ⋅ 8 and concatenating all samples allows us to get a feature matrix. To investigate the proximity of cut-sites the dimensionality of the feature matrix was reduced to two dimensions using UMAP.

To remove the effect of neutrophil elastase, clusters enclosed by valine, isoleucine, alanine and threonine were removed from all samples. The remaining clusters were filtered for each sample separately, to remove clusters which had valine, isoleucine, alanine and threonine as the most common terminals. The cut-site specificities were thereafter calculated as per above.

### Implementation & software

The complete bioinformatic analysis was conducted in Python 3.9. NetworkX and iGraph were used to generate networks. The Leiden algorithm was used as implemented in the leidenalg-package. XGBoost, scikit-learn and SHAP were used for machine learning applications. The UMAP-learn package was used for dimensionality reduction using UMAP. Scipy was used to calculate p-values. The in-house processing package DPKS was used for quantification (39). Vector manipulations utilized numpy. A package for the creation of peptide clusters is available under an MIT license at GitHub: https://github.com/ErikHartman/pepnets. All figures were created in BioRender.com.

### Ethics statement

All animal experiments are performed according to Swedish Animal Welfare Act SFS 1988:534 and were approved by the Animal Ethics Committee of Malmö/Lund, Sweden (permit number M131-16). The use of human wound materials was approved by the Swedish ethical review authority (etikprövningsmyndigheten application number 2023-05051-02). Written consent was received from all subjects prior to participation.

### Pig wound model

In a previous study exploring the effects of the thrombin derived antimicrobial peptide TCP-25, partial thickness wounds of Göttingen minipigs were either infected with *S. aureus* or *P. aeruginosa* (24). The wounds were then dressed with a polyurethane dressing, which was changed 24 and 48 hours post infection, and the old dressings were collected for wound fluid extraction. The same dressing procedure was also performed for uninfected control wounds as well as superinfected wounds which were first infected with *S. aureus*, and infected with *P. aeruginosa* after 24 hours, during the first change of polyurethane dressing. The dressing of the superinfected wound was also collected 72 hours post *S. aureus* infection (24). The collected dressings were placed in syringes and soaked in 10 mM Tris (pH 7.4), before ejecting the fluids. Halt Protease Inhibitor Cocktail (Thermo Fisher Scientific, USA) was then added to half of the extracted wound fluid from each sample. All samples were then stored at -80°C before use.

### Quantitative bacterial counts

The swabs and dressing fluid samples were diluted with sterile PBS to generate 7 10-fold serial dilutions (from 10x to 10^7^x). Six separate 10 L drops of the undiluted sample and each of the dilutions were deposited on a Todd-Hewitt agar plate. The plates were incubated at 37°C in 5% CO2 overnight. The next morning, the number of colonies was counted and recorded.

### Wound fluids from non-healing human wounds

Wound fluid from patients with venous non-healing wounds was collected from Mepilex dressings applied on the wounds for 48-72 hours. The dressings were extracted as described above, and Halt Protease Inhibitor Cocktail added as above before storage at -80°C.

### Identification of bacteria

Colonies from wound swab samples were prepared using the extended direct transfer sample preparation procedure on stainless steel MALDI target plates as described by the manufacturer (Bruker Daltronik GmbH). A Microflex LT/SH SMART MALDI-TOF mass spectrometry (MS) instrument with flexControl v. 3.4 (Bruker Daltronik GmbH) was used to analyze the target plate and collect mass spectra in linear mode over a mass range of 2 to 20 kDa. A spectrum of 240 summed laser shots was acquired for each sample spot. The spectra were analyzed using a MALDI Biotyper (MBT) Compass v. 4.1 with the MBT Compass Library Revision L (DB-9607, 2020) (Bruker Daltronik GmbH).

### Sample preparation, mass spectrometry and data processing

Wound fluid extracts with supplemented protease inhibitor had their protein concentrations measured using the Pierce BCA Protein Assay Kit (Thermo Fisher Scientific, USA) according to the provided instructions. A volume corresponding to 500 µg of protein for pig wound fluids and 100 µg of protein for human wound fluid was diluted in 10 mM Tris (pH 7.4) to 100 µl, and then further diluted in 300 µl 8 M urea (in 10 mM Tris (pH 7.4) to a final concentration of 6 M urea) supplemented with 0.067% RapiGest SF (Waters, USA) (to a final concentration of 0.05% RapiGest SF). The samples were then incubated for 30 minutes at room temperature. Meanwhile, Microcon - 30 centrifugal filter units were rinsed with 100 µl 6 M urea (in 10 mM Tris (pH 7.4)) by centrifugation for 15 minutes at 10 000 RCF at room temperature (RT). The samples were then loaded onto the filter units and centrifuged for 30 minutes at 10 000 RCF at RT, followed by a final rinse with an additional 100 µl 6 M urea (in 10 mM Tris (pH 7.4)) and 5 minutes of centrifugation at 10 000 RCF at RT. The filtrate was then stored at -20°C until LC-MS/MS analysis.

In total, wound fluids from 47 pig wounds (17 wounds infected with *S. aureus* from 4 pigs, 17 wounds infected with *P. aeruginosa* from 4 pigs, 13 uninfected wounds from 4 pigs) at two time points (24 and 48 hours post infection) had their peptides extracted. In addition, wound fluids from 4 superinfected wounds from 1 pig at three different time points (24, 48 and 72 hours post *S. aureus* infection) also had their peptides extracted. 18 human wound fluids from 4 subjects had their peptides extracted.

Extracted peptide samples were acidified by adding 1 µl 100% formic acid (FA) to 60 µl of peptide filtrate. Meanwhile, UltraMicro Spin Columns (The Nest Group, USA) were wet by adding 100 µl 100% acetonitrile (ACN) + 0.1% FA and centrifuging the column at 800 RCF for 1 minute at room temperature. These conditions were used for the remainder of the centrifugation steps of the solid phase extraction. The columns were then equilibrated by centrifuging 100 µl 2% ACN + 0.1% trifluoroacetic acid (TFA) through them twice. Samples were then spun onto the columns, followed by a washing step where 100 µl 2% ACN + 0.1% TFA was centrifuged through. The samples were then eluted by centrifuging 100 µl 70% ACN + 0.1% TFA through the columns. Once eluted, the samples were dried using an Eppendorf Concentrator plus at 45°C and redissolved in 30 µl 2% ACN + 0.1% TFA.

The redissolved peptide samples were loaded onto Evotip Pure columns according to the provided instructions, apart from that the loaded samples were dissolved in 30 µl 2% ACN + 0.1 % FA instead of 20 µl 0.1% FA. These were then analyzed by LC-MS/MS on an Evosep One LC (Evosep, Denmark) coupled to a timsTOF Pro mass spectrometer (Bruker, USA). The LC was equipped with an EV1137 Performance Column – 15 cm x 150 µm, filled with 1.5 µm ReproSil-Pur C18 beads (Evosep, Denmark), and separation was performed using the accompanying 30 samples per day program. The MS used the DDA PASEF mode, doing 10 PASEF scans every acquisition cycle. The accumulation and ramp times were both set to 100 ms. Precursors with a +1 charge were ignored and the target intensity was set to 20 000, with dynamic exclusion active, at 0.4 min. The isolation width was set to 2 at 700 Th and 3 at 800 Th.

The data from the LC-MS/MS runs were searched with PEAKS X. UniProtKB reviewed (Swiss-Prot) protein list of pig proteins was used as a database when searching the pig samples, with the exchange of fibrinogen alpha chain (FIBA_PIG) and fibrinogen beta chain (FIBB_PIG) to the UniProtKB unreviewed (TrEMBL) versions F1RX36_PIG and F1RX37_PIG respectively. When searching the human samples, UniprotKB reviewed (Swiss-Prot) protein list of human proteins was used as a database. The lists can be found in supplemental data X. The precursor tolerance was set to 20 ppm and the fragment tolerance was set to 0.03 Da. Oxidation (M, +15.99) was treated as a possible modification, and a maximum of one modification per peptide was allowed. The search results were filtered at 1% FDR and at least 1 unique peptide for each protein.

The peptide intensities were *log*_2_-transformed. Identified but unquantifiable peptides were imputed by sampling from a uniform distribution *U*(2, 8) which is lower than the least abundant quantifiable peptides. The intensities were then mean-normalized so that all samples had equal intensity-means.

### Zymograms

Zymogram gels were created with a separation gel consisting of 375 mM Tris buffer (pH 8.8), 0.1% (w/v) SDS, 0.1% (w/v) gelatine, 10% (w/v) acryl amide, 0.05% (w/v) TEMED and 0.05% (w/v) APS in Milli-Q water and a stacking gel consisting of 125 mM Tris (pH 6.8), 0.1% (w/v) SDS, 4% (w/v) acryl amide, 0.1% (w/v) TEMED and 0.05% (w/v) APS in Milli-Q water. For each sample, 5 µg of protein without added protease inhibitor was diluted to 5 µl with Milli-Q water, and then mixed with 5 µl sample buffer consisting of 400 mM Tris-HCl (pH 6.8), 20% (v/v) glycerol, 5% (w/v) SDS and 0.03% (w/v) bromophenol blue in Milli-Q water, which was then added to the wells. The gels were then run using an electrophoresis buffer consisting of 25 mM Tris, 200 mM glycine and 0.1% (w/v) SDS in Milli-Q water at pH 8.7 for 60 minutes at 150 V. Afterwards, the gels were washed with deionized water and incubated for 60 minutes in 2.5 % Triton X-100 at room temperature, with 160 rpm shaking, and followed by another deionized water wash. Next, the gels were incubated overnight at 37°C in an enzyme buffer consisting of 50 mM Tris-HCl (pH 7.5), 200 mM NaCl, 5 mM CaCl2 and 1 µM ZnCl2 with 50 rpm shaking. The next day, the gels were washed in deionized water and incubated in a staining buffer consisting of 0.25% (w/v) Coomassie brilliant Blue G-250, 38.4% (v/v) ethanol and 7% (v/v) acetic acid in Milli-Q water for 60 minutes. The gels were then placed in a de-staining solution consisting of 9.6% (v/v) ethanol and 7% (v/v) acetic acid in Milli-Q and imaged using a Chemidoc MP Imaging System (Bio-Rad Laboratories, USA).

### SDS-PAGE

From each wound fluid sample, 20 µg of protein with added protease inhibitor was mixed with Milli-Q water to 8 µl. 10 µl Tricine SDS Sample Buffer (2X) and 2 µl NuPAGE Reducing Agent (10X) was added to each sample. The samples were then incubated at 95 °C for 5 minutes. 10-20% Tricine gels and running buffer were prepared as described by the manufacturer’s instructions and were then run for 90 minutes at 100 V. Once the runs were finished, the gels were stained with Gelcode Blue Safe Protein Stain (Thermo Fisher Scientific, USA) according to the instructions provided by the manufacturer. Imaging was then performed using a Chemidoc MP Imaging System (Bio-Rad Laboratories, USA).

## Code availability

A package containing the code used for the creation and analysis of peptide clusters is available in the open GitHub repository https://github.com/ErikHartman/pepnets under an MIT license.

## Data availability

The data in this study will be made available at ProteomeXchange at the time of publication.

## Acknowledgement

We thank the Swedish National Infrastructure for Biological Mass Spectrometry (BioMS) for performing the LC-MS/MS analysis. Xinnate AB provided the funding and project management resources enabling the clinical safety study on non-healing wounds that generated the control samples used in this work. We are indebted to Drs. Sigrid Lundgren, Karl Wallblom, and colleagues at the Department of Dermatology and the Clinical Trial Unit at Skane University Hospital Lund for patient related work and for providing the patient wound fluids used in this study, and to Dr. Bo Nilson, Division of Medical Microbiology, Department of Laboratory Medicine, Lund University, for identification of bacteria isolated from non-healing ulcers. We acknowledge support by grants from the Swedish Research Council (project 2017-02341, 2020-02016), Edvard Welanders Stiftelse and Finsenstiftelsen (Hudfonden), the Royal Physiographic Society, the Crafoord and Österlund Foundations, and the Swedish Government Funds for Clinical Research (ALF).

## Supplementary information

### Supplementary Notes S1. Characterisation of the porcine wound fluid peptidomes

The infected wounds developed clinical signs of infection, including erythema and visible bacterial biofilms (Supplementary Fig. 1a). The bacterial load in the single infections remained stable with around 10^3^ CFU (Supplementary Fig. 1b). In the double infections, however, the CFU count of *S. aureus* increased when introducing *P. aeruginosa*, and the two species reached similar levels of colonization on day 3 (Supplementary Fig. 1c). Dimensionality reduction using uniform manifold projection (UMAP) on the log-transformed peptide intensities showed that peptidomes are sample type-specific (Supplementary Fig. 1d). The number of identified peptides decreased between days *n* and *n* + 1, with an average decrease of 34% (Supplementary Fig. 1e). Hierarchical clustering on peptide intensities revealed two distinct clusters, where one cluster constitutes the shared peptides, and the second contains most peptides identified in a subset of samples (Supplementary Fig. 1f). Peptide lengths vary between 7 and 58 amino acids, with a mean length of 15.2 amino acids. Control samples contain a smaller fraction of short peptides (<11 amino acids) compared to infected samples, and peptide length decreases between days 1 and 2 (Supplementary Fig. 2d).

### Supplementary Notes S2. Peptide properties in clusters

Previous work has shown that peptide properties are largely conserved in peptides with high sequence similarity (1,2). To investigate this and to motivate the use of peptide clusters as a functional entity, the antimicrobial tendencies of all peptides were predicted using a pre-trained deep convolutional neural network (3). The network outputs a classification probability (antimicrobial prediction score) for a given peptide sequence. The variance of the antimicrobial prediction score between peptides within a cluster was compared to the variance between all peptides in a protein as a benchmark. The inter-cluster variance is significantly lower than the inter-protein variance of antimicrobial scores (Supplementary Fig. 2). Other properties, such as isoelectric point and aromaticity, showed similar distributions.

### Supplementary Notes S3. Biological significance of HMGB1, HPT and PR39

High mobility group-box 1 (HMGB1) is a dual-functioning protein, as its intracellular role entails transcriptional regulation by binding to DNA while it acts as a damage-associated molecular pattern (DAMP) when released into the extracellular environment. It contains three functional regions denoted as the A-box, B-box and the acidic C-terminal tail (Fig 4a). Further, it contains heparin-binding, TLR-binding and RAGE-binding domains which regulate the inflammatory response. The degradation of HMGB1 into functional peptides by endogenous proteases have been studied, demonstrating cleavage by C1, neutrophil elastase, cathepsin G and matrix metalloproteinase 3 (32–34). HMGB1 has also been documented to interact with *P. aeruginosa* in pneumonia and cystic fibrosis by mediating inflammation and attenuating bacterial clearance (35), as well as in keratitis (36). We find that cleavage of HMGB1 in wounds infected by *P. aeruginosa* produced peptides from the A-box region (29–43) and B-box (112–127) as well as the acidic tail (190–206, 196–212) which were not identified in*S. aureus*infections or control. Additionally, these clusters appeared in double infected wounds on day 3, when the bacterial load of *P. aeruginosa* had increased. The two clusters from the A-box and B-box are surrounded by cut sites with lysine at p1, matching the cut site specificity of P. aeruginosa.

Haptoglobin (HPT) is a protein that binds free hemoglobin, thereby protecting tissues from deleterious oxidative activity. However, it has also been demonstrated to bind to HMGB1, and the resulting complex binds CD163, leading to anti-inflammatory signaling in the monocyte-macrophage lineage (37). In *P. aeruginosa* infections, HPT is cleaved in the region 100-120 resulting in differentially abundant peptide clusters although also present in S. aureus-infected wounds (Fig. 2d). However, the peptides from this region are subsequentially cleaved at a cut site with a lysine at position p1, into two sub-peptides (Fig. 4c). These are only identified in *P. aeruginosa*-infected wounds. Additionally, the peptide cluster in region 154-169 is only identified in *P. aeruginosa*-infected wounds and is flanked by cut sites with a lysine at p1.

PR-39 is an antibacterial protein, with peptides derived from the region of PR39 (131–169) exhibiting high antibacterial activity. Cleavage in this region has been documented to inhibit the antimicrobial effect, as the shorter sub-peptides are much less potent than the full peptide (27). All clusters of PR-39 exhibit similar abundance, except for clusters in this region, which are more abundant in *P. aeruginosa*-infected wounds. Both *P. aeruginosa* and*S. aureus*has been demonstrated to exhibit resistance towards the effect of PR-39 -*S. aureus*being completely resistant to high concentrations and *P. aeruginosa* being relatively resistant to PR-AMPs.

### Supplementary Notes S4. Description of human non-healing wound samples

Out of the 18 samples, 10 showed growth of *P. aeruginosa*. These samples also often contained other bacteria, such as *E. faecalis*, **S. lugdunensis, Corynebacterium sp., S. aureus*, and* S. agalactiae*. The other 8 samples primarily contained* S. aureus*. Sample 9 contained* P. aeruginosa* but did not show a *P. aerguinosa*-like profile on the zymogram, and is therefore labeled “P. aeruginosa MALDI+ Zymo-”. One patient was given antibiotics against a *P. aeruginosa*-infection, and two samples derived from this patient are therefore labeled as “P. aeruginosa after antibiotics”.

## Supplementary figures

**Supplementary Fig. 1.**
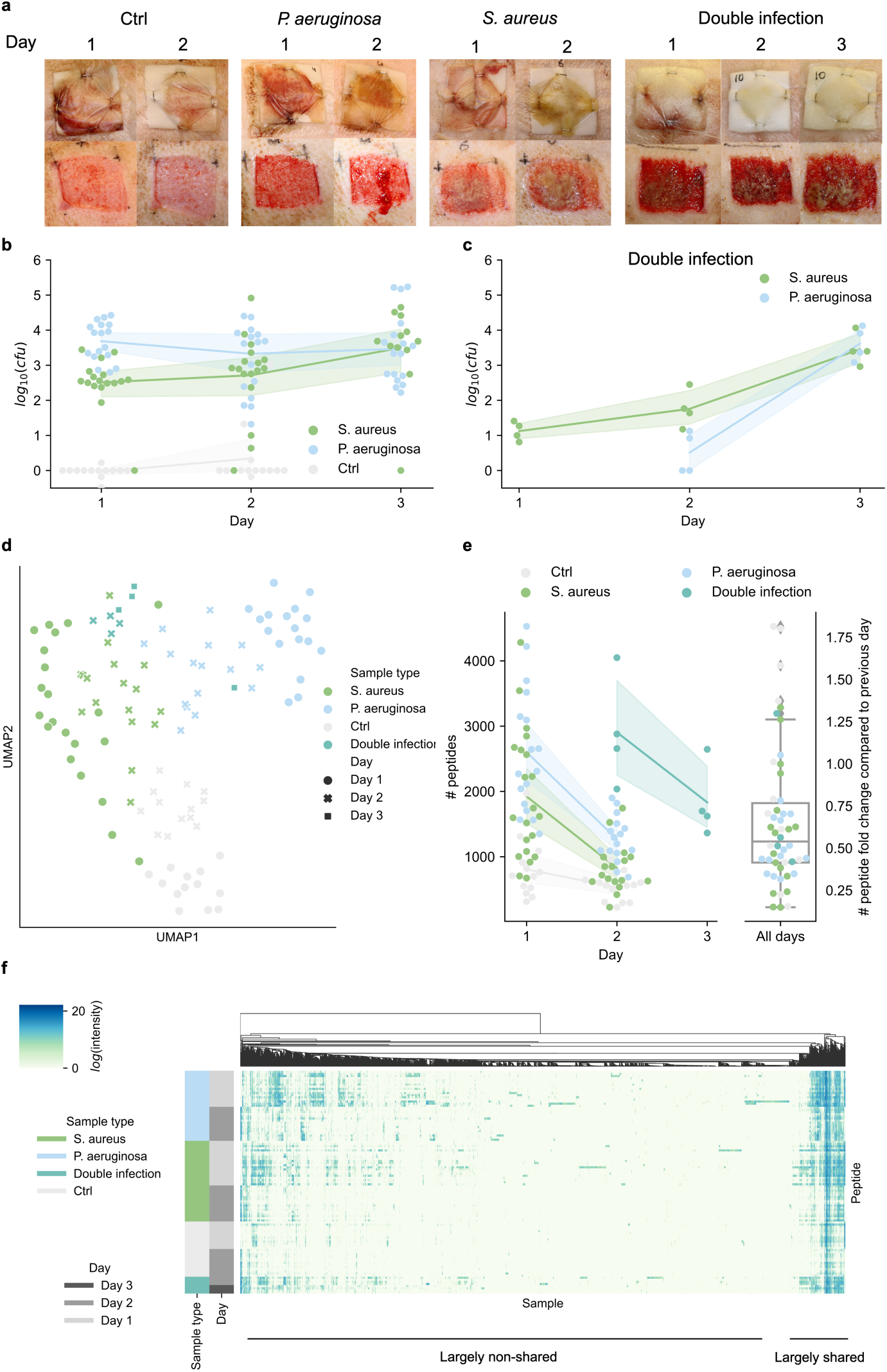
The wound fluid peptidome is pathogen-specific. **a** Representative pictures of the porcine wounds with and without dressing over the sampling period. Wounds were infected on day 0. The double infected wounds were infected with *P. aeruginosa* on day 1. **b** *log*_10_CFU over the three-day period in the single infections and control wounds. **C** *log*_10_CFU over the three-day period in the double infected wounds. In **b** and **c** the line shows the mean trend and the error bars ±1SD. **d** Uniform manifold projection (UMAP) of the peptide intensities colored by sample type and shaped determined by the day the sample was taken. **e** Number of peptides detected over the timespan. The right panel shows the fold change of number of peptides between day *n* and *n* − 1. Every sample is shown as a scatter, alongside a line for mean values. For the means, ±1 SD is shown as error-bands. **f** Clustering of peptide groups using hierarchical clustering. The color in the heatmap is proportional to the normalized *log*_2_-transformed intensity. Clustering is performed using hierarchical clustering minimizing the Ward-distance.

**Supplementary Fig. 2.**
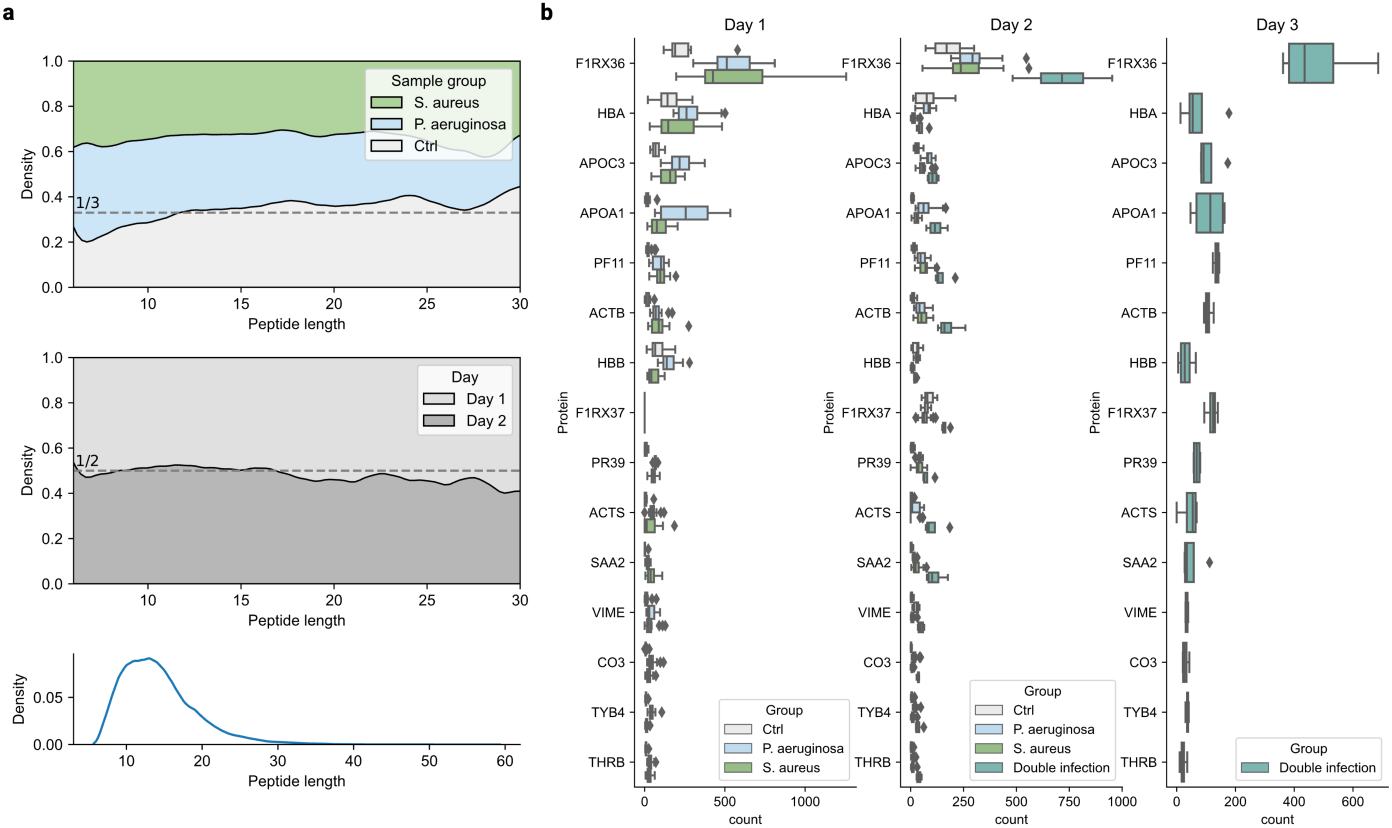
Peptide characteristics. **a** Length distribution of peptides split on sample type (upper) and day (middle). The lower panel shows the kernel density estimate of the length of all peptides. **e** Number of peptides mapped to different proteins. Top 15 proteins are shown.

**Supplementary Fig. 3.**
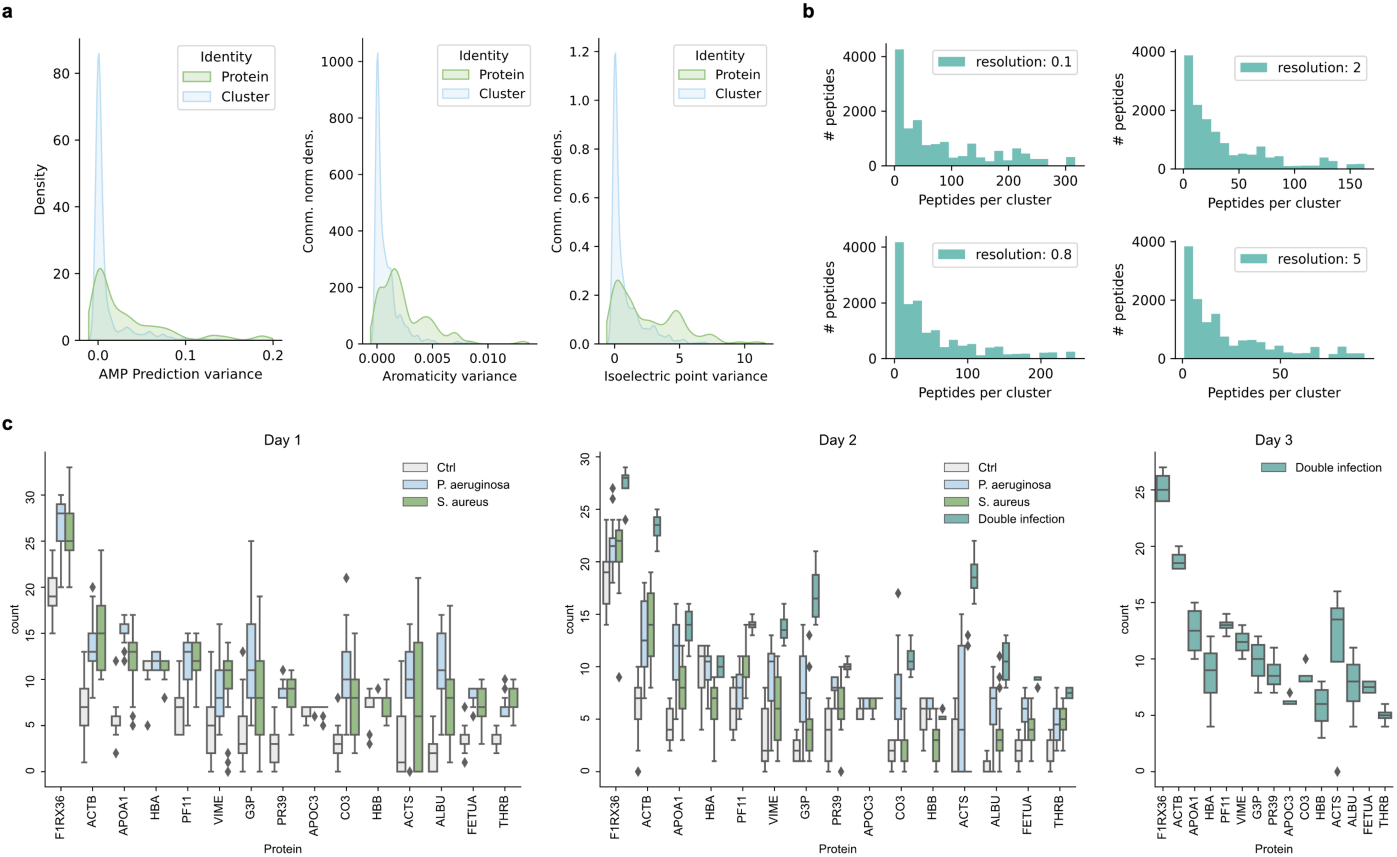
Cluster characteristics. **a** Inter-cluster and inter-protein variance of aromaticity and isoelectric point of peptides. **b** Cluster size distribution when having different values for the *Γ*-parameter (resolution) in the Leiden algorithm. **c** Number of clusters mapped to different proteins.

**Supplementary Fig. 4.**
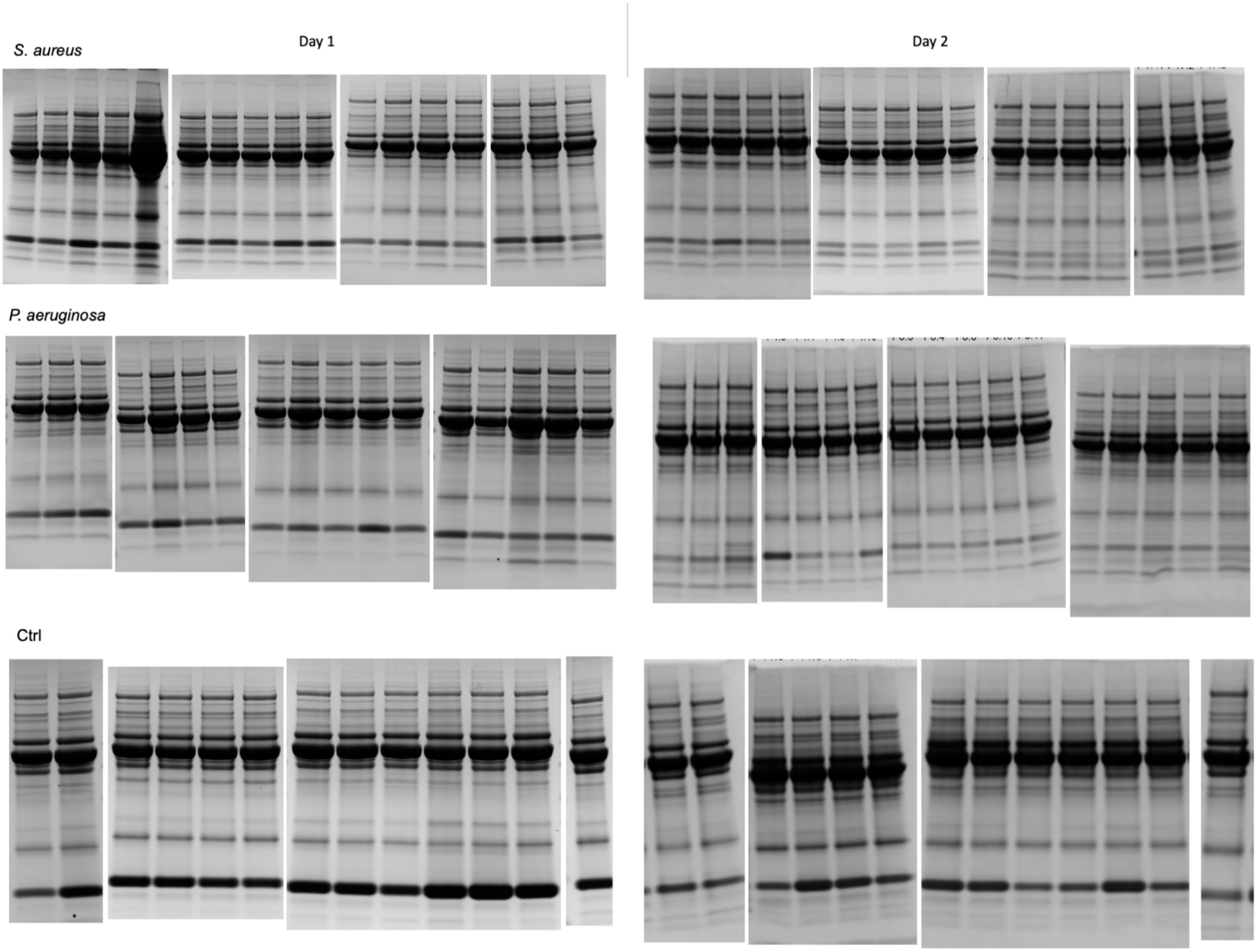
SDS-PAGE for sample types. SDS-PAGE on the wound fluid of all samples on day 1 (left) and day 2 (right). The upper row shows the sample s infected by *S. aureus*, the middle by *P. aeruginosa* and the lower the uninfected control samples.

**Supplementary Fig. 5.**
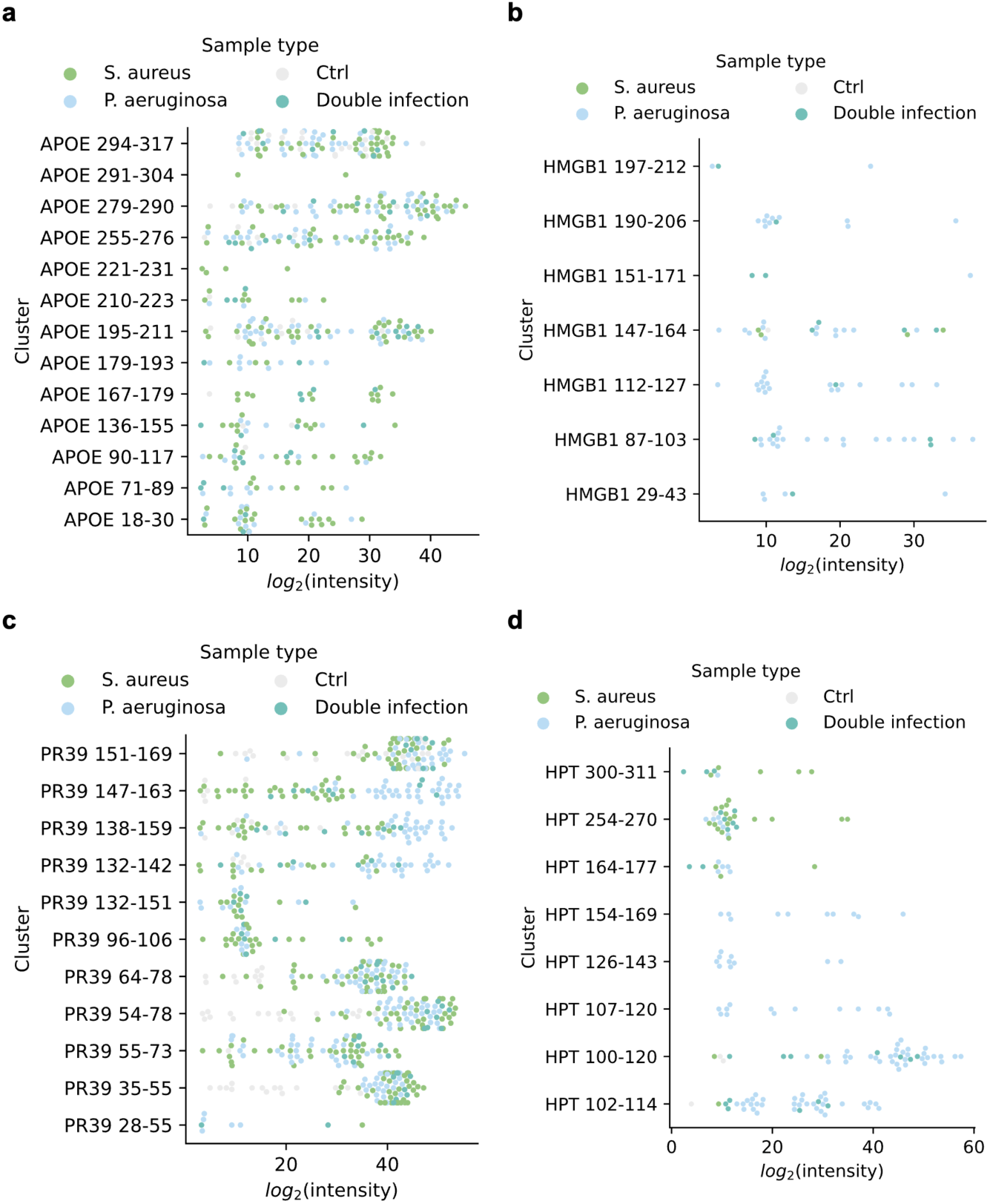
Cluster abundances. Abundances for all clusters in **a** APOE, **b** HMGB1 **c** PR39 and **d** HPT.

**Supplementary Fig. 6.**
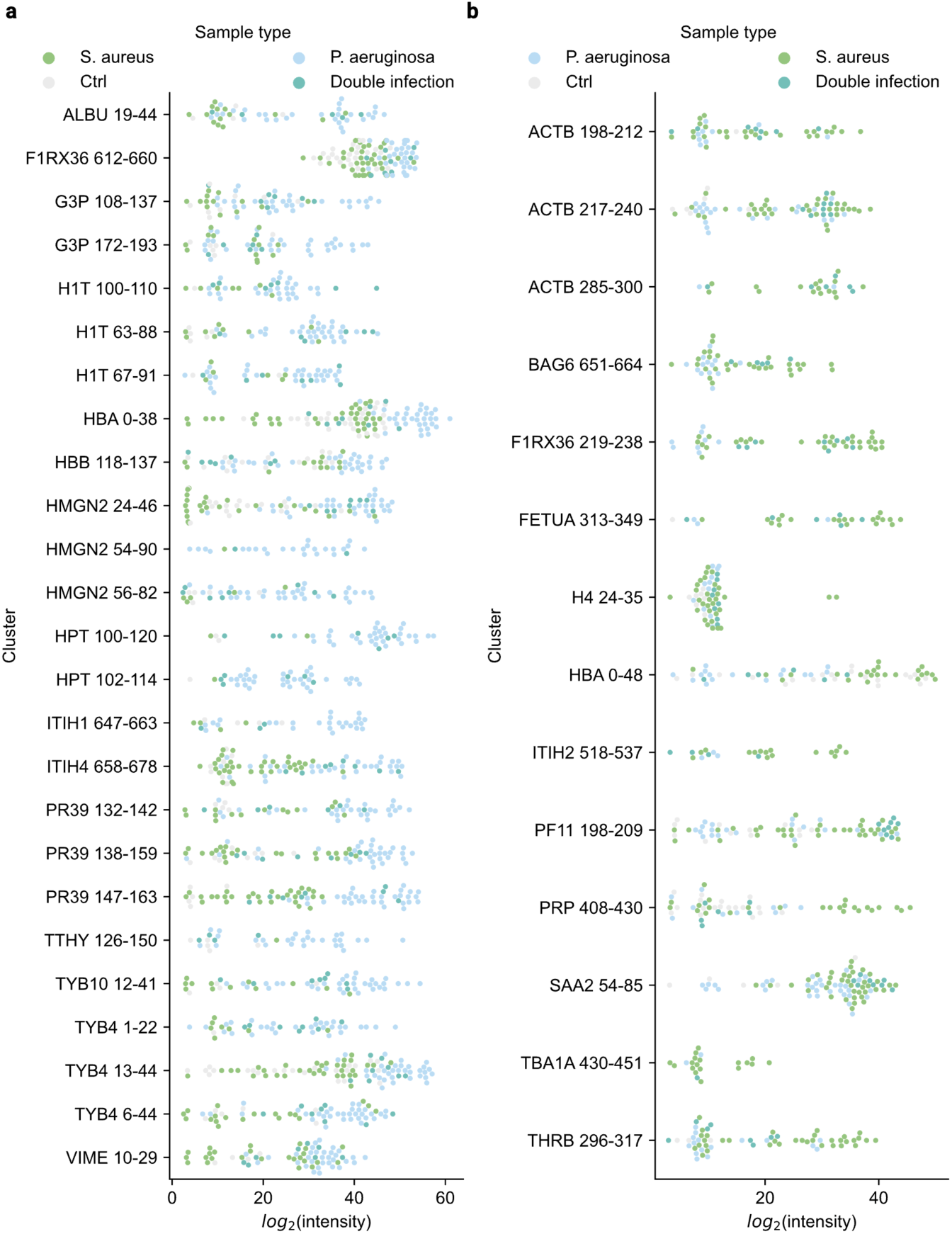
Top 50 differentially expressed cluster abundances. Abundances of the clusters exhibiting the highest differential expression in **a** *P. aeruginosa*-infected wounds and **b** *S. aureus*-infected wounds.

**Supplementary Fig. 7.**
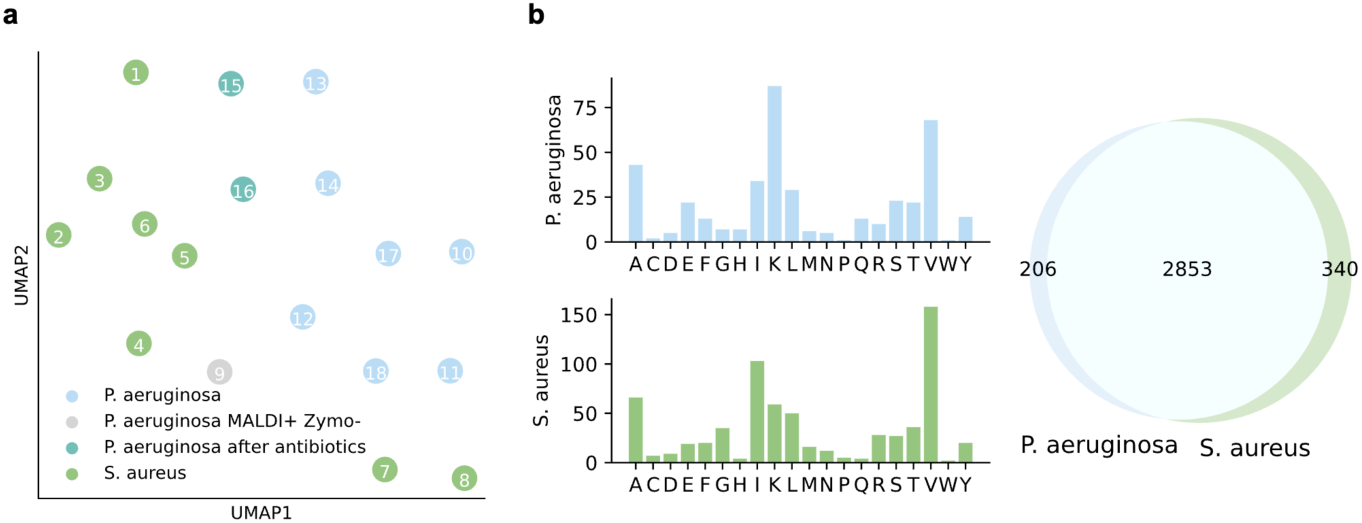
a. UMAP of the quantified peptide clusters colored based on results from MALDI and zymograms. **b** Amino acid profile for the amino acids at the p1 position for the unique clusters when stratifying the samples on the species of the primary colonizer.

**Supplementary Fig. 8.**
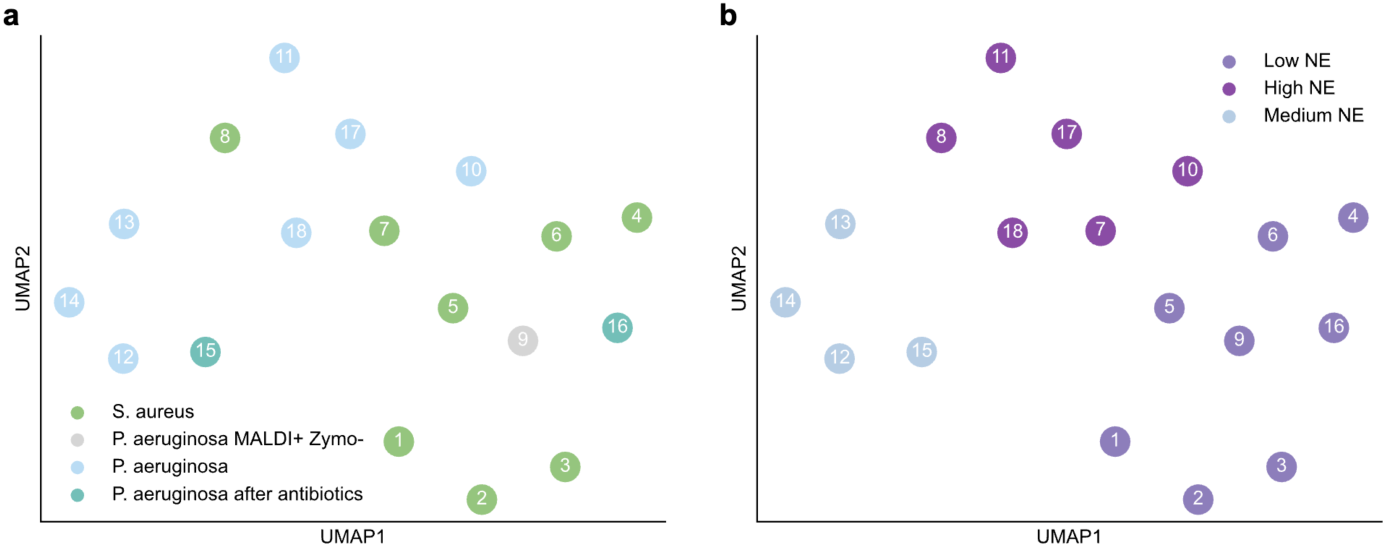
Removing the effect of neutrophil elastase. The effect of neutrophil elastase was removed by removing all clusters which had valine, isoleucine, alanine and threonine at the p1 position of any terminal. **a,b** UMAP projection of the p4-p4’ KL-weighted amino acid frequencies. In **a** the scatters are colored depending on the appearance of the zymogram and the bacterial composition of the MALDI. In **b** the scatters are colored based on the identified groups of high, medium and low NE-like p1 specificity.

**Supplementary Table 1.** The 50 top ranking features extracted by training an XGBoost classifier and estimating the feature importance using SHAP. The topn-columns contain the sum of the abundance of the cluster per group calculated by taking the sum of the top 3 most abundant peptides.

| Rank | Protein | Start | End | Ctrl<br>topn | *P. aeruginosa*<br>topn | *S. aureus*<br>topn |
| --- | --- | --- | --- | --- | --- | --- |
| 0 | HPT | 102 | 114 | 0.15 | 25.4 | 0.25 |
| 1 | PF11 | 48 | 62 | 4.44 | 36.34 | 32.76 |
| 2 | SAA2 | 54 | 85 | 3.37 | 27.72 | 34.25 |
| 3 | HMGN2 | 54 | 90 | 0 | 18.51 | 0.3 |
| 4 | APOC3 | 39 | 83 | 40.88 | 51.65 | 50.34 |
| 5 | HBA | 0 | 22 | 40.1 | 45.7 | 34.2 |
| 6 | F1RX37 | 48 | 73 | 20.44 | 23.4 | 17.65 |
| 7 | THRB | 296 | 317 | 0.83 | 6.49 | 21.55 |
| 8 | RL31 | 90 | 106 | 1.27 | 3.5 | 5.35 |
| 9 | PF11 | 12 | 39 | 10.9 | 39.65 | 40.86 |
| 10 | ACTB | 227 | 240 | 0.72 | 6.14 | 12.77 |
| 11 | CO3 | 437 | 457 | 0 | 18.79 | 0 |
| 12 | F1RX36 | 219 | 238 | 0 | 1.92 | 23.08 |
| 13 | PF11 | 198 | 209 | 7.18 | 12.04 | 25.8 |
| 14 | G3P | 192 | 213 | 2.71 | 15.05 | 0.5 |
| 15 | HMGB2 | 192 | 210 | 0 | 20.69 | 0 |
| 16 | F1RX36 | 593 | 623 | 24.64 | 39.74 | 43.33 |
| 17 | F1RX37 | 167 | 186 | 9.66 | 6.2 | 4.43 |
| 18 | TBB5 | 422 | 444 | 1.48 | 9.37 | 10.87 |
| 19 | G3P | 192 | 217 | 16.56 | 16.37 | 16.31 |
| 20 | PF11 | 128 | 161 | 24.55 | 36.9 | 49.2 |
| 21 | APOE | 255 | 276 | 1.96 | 16.46 | 16.2 |
| 22 | BAG6 | 651 | 664 | 0 | 3.99 | 13.1 |
| 23 | F1RX37 | 48 | 87 | 16.97 | 20.11 | 17.92 |
| 24 | APOE | 294 | 317 | 19.08 | 19.43 | 25.48 |
| 25 | VTNC | 65 | 77 | 0 | 1.04 | 3.72 |
| 26 | ITIH4 | 632 | 648 | 2.4 | 2.45 | 6.72 |
| 27 | HBB | 29 | 61 | 26.81 | 21.87 | 6.11 |
| 28 | HBB | 118 | 137 | 12.66 | 32.71 | 8.71 |
| 29 | HBA | 0 | 48 | 24.32 | 8.09 | 22.99 |
| 31 | F1RX37 | 55 | 87 | 23.12 | 22.67 | 21.93 |
| 32 | TTHY | 126 | 150 | 1.8 | 23.67 | 0.6 |
| 33 | LOX15 | 624 | 643 | 2.24 | 1.82 | 6.06 |
| 34 | ANXA1 | 8 | 29 | 14.35 | 28.95 | 20.64 |
| 35 | HMGN2 | 24 | 46 | 10.7 | 36.88 | 9.21 |
| 36 | PF11 | 161 | 203 | 2.23 | 4.71 | 10.37 |
| 37 | TYB4 | 13 | 44 | 24.27 | 48.98 | 27.38 |
| 38 | VIME | 47 | 71 | 4.04 | 26.85 | 15.77 |
| 39 | PF11 | 171 | 198 | 0.87 | 5.28 | 6.49 |
| 40 | APOC3 | 20 | 43 | 33.62 | 48.21 | 41.66 |
| 41 | ITIH4 | 658 | 678 | 3.4 | 34.59 | 16.44 |
| 42 | VIME | 38 | 68 | 5.91 | 10.28 | 9.73 |
| 43 | H4 | 24 | 35 | 2.27 | 3.41 | 9.34 |
| 44 | SAA2 | 84 | 119 | 2.7 | 20.42 | 17.4 |
| 45 | F1RX36 | 466 | 518 | 36.85 | 45.09 | 44.81 |
| 46 | ITIH2 | 492 | 521 | 0 | 5.94 | 9.07 |
| 47 | PF11 | 137 | 166 | 40.68 | 39.57 | 48.2 |
| 48 | FA5 | 985 | 1009 | 1.34 | 9.09 | 2.78 |
| 49 | VTNC | 108 | 127 | 0 | 0.27 | 2.23 |
| 50 | HMGB2 | 1 | 18 | 0.44 | 3.21 | 4.22 |

**Supplementary Table 2.**
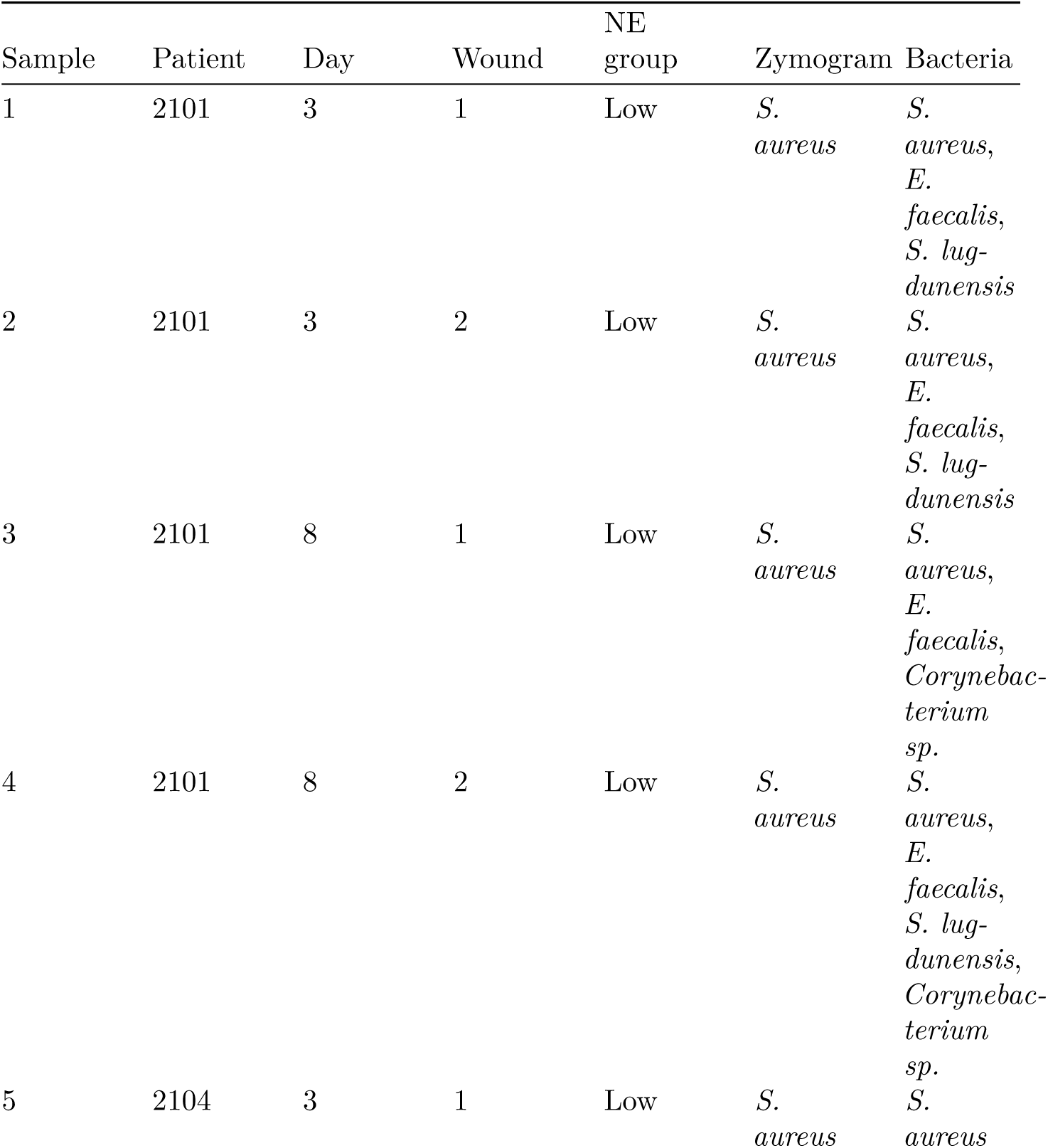
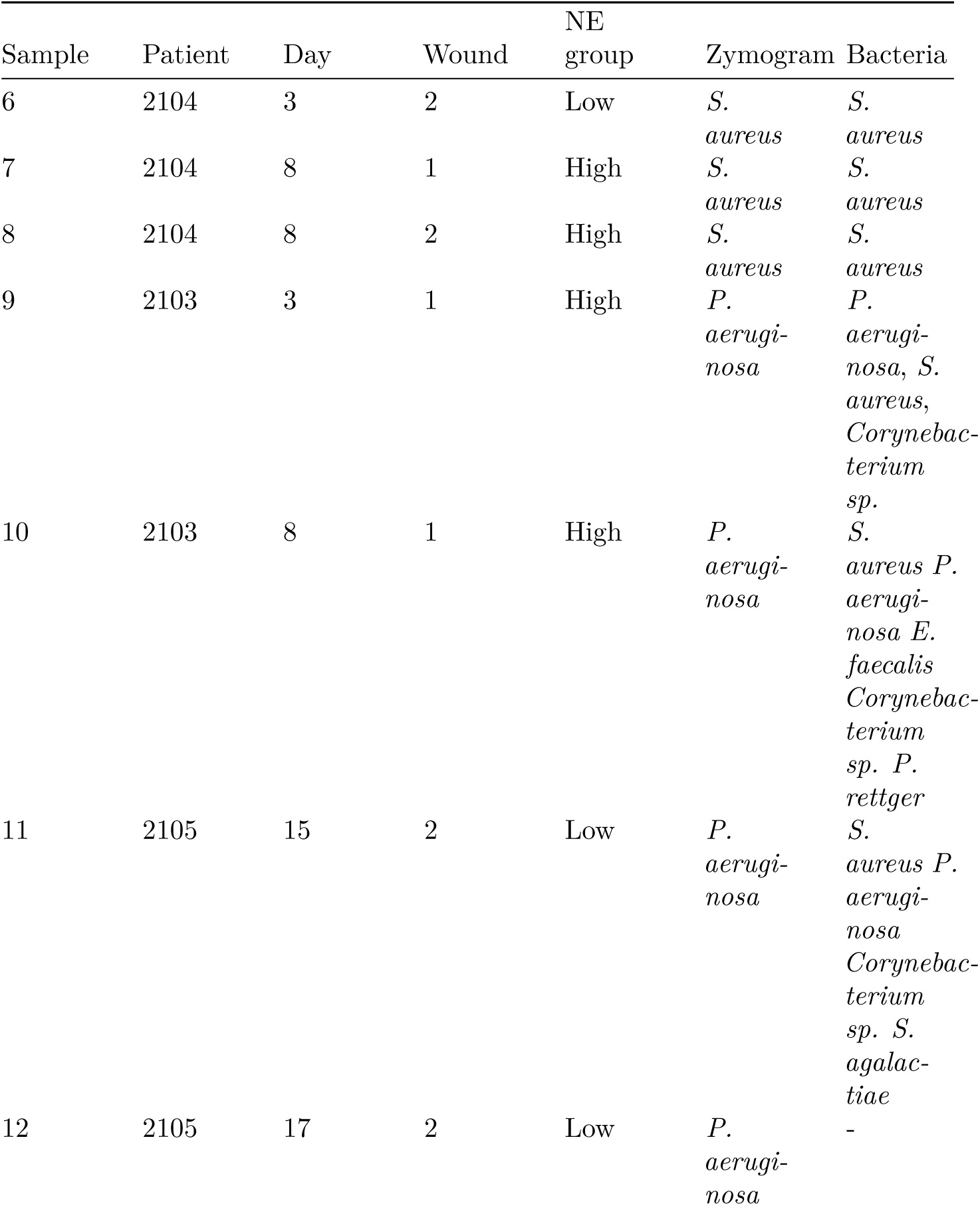
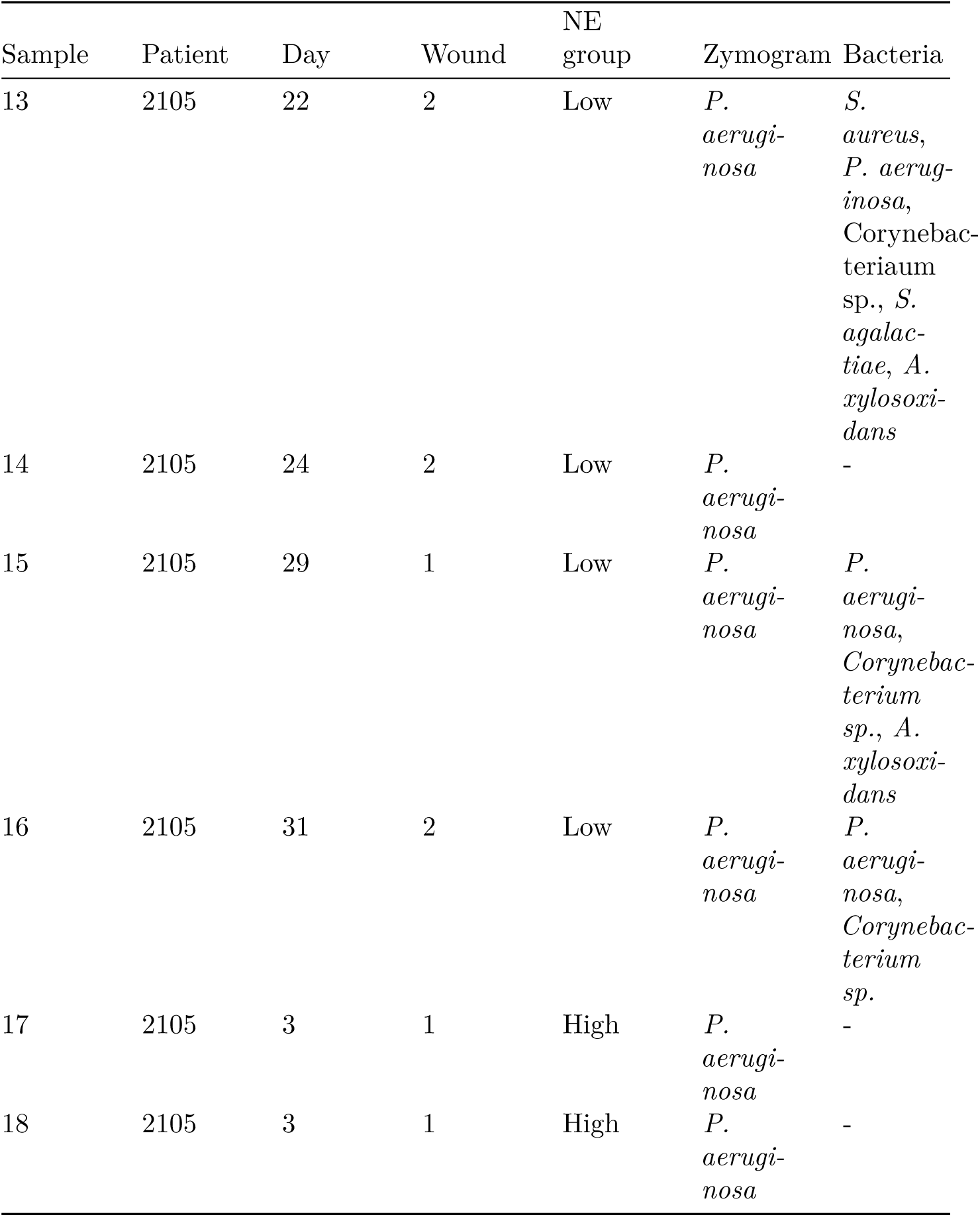
Sample specifications for human samples. Bacterial identification by MALDI was not performed on samples 17 and 18.

| Sample | Patient | Day | Wound | NE<br>group | Zymogram | Bacteria |
| --- | --- | --- | --- | --- | --- | --- |
| 1 | 2101 | 3 | 1 | Low | *S. aureus* | *S. aureus*,<br>*E. faecalis*,<br>*S. lug-*<br>*dunensis* |
| 2 | 2101 | 3 | 2 | Low | *S. aureus* | *S. aureus*,<br>*E. faecalis*,<br>*S. lug-*<br>*dunensis* |
| 3 | 2101 | 8 | 1 | Low | *S. aureus* | *S. aureus*,<br>*E. faecalis*,<br>*Corynebac-*<br>*terium*<br>*sp.* |
| 4 | 2101 | 8 | 2 | Low | *S. aureus* | *S. aureus*,<br>*E. faecalis*,<br>*S. lug-*<br>*dunensis*,<br>*Corynebac-*<br>*terium*<br>*sp.* |
| 5 | 2104 | 3 | 1 | Low | *S. aureus* | *S. aureus* |
| 6 | 2104 | 3 | 2 | Low | *S.*<br>*aureus* | *S.*<br>*aureus* |
| 7 | 2104 | 8 | 1 | High | *S.*<br>*aureus* | *S.*<br>*aureus* |
| 8 | 2104 | 8 | 2 | High | *S.*<br>*aureus* | *S.*<br>*aureus* |
| 9 | 2103 | 3 | 1 | High | *P.*<br>*aerugi-*<br>*nosa* | *P.*<br>*aerugi-*<br>*nosa*, *S.*<br>*aureus*,<br>*Corynebac-*<br>*terium*<br>*sp.* |
| 10 | 2103 | 8 | 1 | High | *P.*<br>*aerugi-*<br>*nosa* | *S.*<br>*aureus* *P.*<br>*aerugi-*<br>*nosa* *E.*<br>*faecalis*<br>*Corynebac-*<br>*terium*<br>*sp.* *P.*<br>*rettger* |
| 11 | 2105 | 15 | 2 | Low | *P.*<br>*aerugi-*<br>*nosa* | *S.*<br>*aureus* *P.*<br>*aerugi-*<br>*nosa*<br>*Corynebac-*<br>*terium*<br>*sp.* *S.*<br>*agalac-*<br>*tiae* |
| 12 | 2105 | 17 | 2 | Low | *P.*<br>*aerugi-*<br>*nosa* | - |
| 13 | 2105 | 22 | 2 | Low | *P. aeruginosa* | *S. aureus*,<br>*P. aeruginosa*,<br>*Corynebacterium*<br>*sp.*, *S. agalactiae*, *A. xylosoxidans* |
| 14 | 2105 | 24 | 2 | Low | *P. aeruginosa* | - |
| 15 | 2105 | 29 | 1 | Low | *P. aeruginosa* | *P. aeruginosa*,<br>*Corynebacterium*<br>*sp.*, *A. xylosoxidans* |
| 16 | 2105 | 31 | 2 | Low | *P. aeruginosa* | *P. aeruginosa*,<br>*Corynebacterium*<br>*sp.* |
| 17 | 2105 | 3 | 1 | High | *P. aeruginosa* | - |
| 18 | 2105 | 3 | 1 | High | *P. aeruginosa* | - |

